# Ku limits aberrant mRNA splicing promoted by intronic antisense Alu elements

**DOI:** 10.1101/2025.11.20.689478

**Authors:** Giovanni Pascarella, Madison P Turley, Mariia Mikhova, Gargi Parkhi, Jared Godfrey, Joshua Heyza, Tomáš Janovič, Li Han, Kefei Yu, Andrew Olive, Piero Carninci, Jens C Schmidt, Katheryn Meek

## Abstract

Alu elements are short repeats that occupy approximately 10% of the human genome. The enrichment of primate genomes with Alu sequences occurred at the prosimian/new-world monkey evolutionary juncture, which coincided with a substantial increase in the expression levels of the Ku70/80 heterodimer. Previous work suggests that the elevated levels of Ku70/80 are required to prevent the activation of innate immune signaling pathways triggered by RNA molecules derived from Alu elements. Here we demonstrate that Ku regulates mRNA splicing, by inhibiting the use of alternative splice sites contained in intronic antisense Alu elements. Dysregulation of splicing precedes cell death and impacts genes involved in essential RNA metabolism processes, including splicing and ribosome biogenesis, likely impacting cell viability. We propose a model in which Ku binding of antisense Alu elements in introns of mRNAs modulates the use of alternative splice sites to balance beneficial and detrimental contributions of Alu repeats within primate genomes.

## INTRODUCTION

The primary function of the DNA-dependent protein kinase complex is recognition of DNA double-strand breaks (DSBs) during non-homologous end-joining (NHEJ). This complex contains the DNA end-binding heterodimer, Ku70 and Ku80, and the large catalytic subunit, DNA-PKcs [reviewed in ^1^]. In NHEJ, after binding the two DNA ends, DNA-PK synapses the ends DSBs and then anchors the assembly of NHEJ’s ligase complex (Ligase 4, XRCC4, XLF, and PAXX) into several distinct configurations that facilitate end-processing and eventual end ligation ^2–4^. It has been well-established that human cells express dramatically higher levels of all three DNA-PK components than non-primate cells ^5,6^. We previously reported that the increase in expression levels of Ku70, Ku80, and DNA-PKcs occured evolutionarily when new world monkeys diverged from prosimians (primitive primates) ^7^. Remarkably this coincides precisely with the expansion of repetitive Alu elements in the genomes of higher primates ^7^. Alu elements mediate the vast majority of non-allelic homologous recombination (NAHR) in human cells ^8^, likely an important factoid that explains many aspects of primate evolution. But NAHR is often deleterious ^9,10^, and our study suggested that DNA-PK may modestly restrain Alu-mediated NAHR ^7^.

It has been known for over a decade that Ku80 is essential in human cells, but not rodent cells ^11,12^. Although DNA-PKcs is not essential in human cell lines, the absence of patients with loss of function mutations in DNA-PKcs suggests that it is essential for organismal viability in humans ^7,13^. Recently, new functions, potentially independent of DNA end-joining have been ascribed to the Ku heterodimer ^14,15^, and it is likely those functions, not its role in NHEJ, that are essential for human cell viability. Hendrickson and colleagues were the first to establish that Ku80 is essential in cultured human cells ^11,12^; cell death was attributed to telomeric loss by a hyper-recombination mechanism ^16^. The telomere-dependent essentiality of Ku has recently been revisited by Kelly et al. ^14^, and Zhu et al. ^15^. Both studies concluded that telomeric loss did not accurately explain the rapid cell death observed after Ku ablation in human cells. In the work by Kelly et al., the authors could not define a single pathway that was impacted by Ku depletion but proposed that numerous cellular processes were disrupted including RNA related processes, translation, ribosome biogenesis, cell cycle and mitotic regulation. In contrast, Zhu et al. proposed that high levels of Ku are required to bind abundant Alu-encoded RNAs expressed in higher primates, preventing these RNAs from activating innate immune pathways. They suggested that the temporal correlation between the increased expression levels of DNA-PK and the genomic expansion of Alu elements ^7^, was an adaptation to restrain Alu-encoded RNAs from activating innate immunity.

The distribution of Alu elements in the genomes of higher primates is biased towards gene rich regions, with approximately 650,000 Alu elements being present in transcribed regions^17,18^. Many of these Alu element insertions in both the sense and antisense orientation are found in introns, where they can have profound impacts on RNA splicing, due to the presence of cryptic splice sites in the Alu RNA sequence ^19^. Importantly, higher primates have evolved mechanisms to prevent dysregulation of splicing by Alu element insertions into gene bodies. For example, hnRNP C has been shown to directly compete with the key splicing factor U2AF65 for binding to the polypyrimidine tracts upstream of splice acceptor sites within intronic antisense Alu elements to prevent inclusion of Alu exons ^18^. In addition, the RNA helicase DHX9 resolves secondary structures formed by inverted repeat Alu elements pairs (irAlus) within transcripts to prevent aberrant RNA processing ^20^. While mis-splicing of intronic Alu elements must be tightly controlled, their presence has contributed to transcriptome evolution, analogous to the role of Alu elements in NAHR -- both processes shaping the evolution of higher primates.

To define the essential function of the Ku complex in human cells, we leveraged a collection of cells lines that allowed specific degradation of key NHEJ factors using a proteolysis targeting chimera (PROTAC). Contradicting previous work ^15^, we do not observe activation of innate immune signaling pathways in two human cell lines depleted of Ku70. In contrast, we find that degradation of Ku70 results in substantial dysregulation of mRNA splicing. The splicing alterations are conserved across multiple cell lines and impact a core set of genes involved in RNA processing, which likely impacts processes essential for cell survival including ribosome biogenesis and mRNA maturation. The magnitude of the splicing changes observed after Ku function is lost are comparable to the disruption of well-established splicing factors, making the Ku complex a *bona fide* splicing regulator. Mechanistically, we demonstrate that Ku binds to a stem loop within antisense Alu elements to suppress the use of cryptic splice acceptor sites immediately upstream of this conserved structural element. We propose a unified model in which high levels of Ku70/80 in higher primates are required to bind abundant antisense Alu RNA within introns to prevent the use of cryptic splice sites contained in the antisense Alu sequences.

## RESULTS

### Cellular ablation of Ku70, or Ku80 but not XLF or DNA-PKcs results in rapid cell death in U2OS cells

Because there is still much to understand about the precise essential requirement for Ku and DNA-PKcs in higher primates, we took advantage of our recently developed reagents including a panel of U2OS cell lines that were gene edited to incorporate a Flag-HaloTag (referred to as Halo) at the N-terminus to both alleles of XLF, Ku70, or DNA-PKcs and a commercially available proteolysis targeting chimera Halo ligand (Halo-PROTAC) to degrade Halo-tagged proteins in living cells ^21,22^. For this study, we also generated U2OS cells expressing Halo-Ku80 from its endogenous locus (Fig. S1A,B). The Halo-DNA-PKcs had a moderate cell fitness deficit while the Halo-Ku80 cell line a more substantial reduction in cell growth (as evidenced by small colony size, Fig. S1C), and the Halo-DNA-PKcs cell strain had a mild NHEJ deficiency (Fig. S1D). The Halo-PROTAC efficiently depleted the NHEJ factors as assessed by staining with a fluorescent Halo ligand (JFX650) and western blotting (Fig. 1A,B). Depletion of either Halo-Ku70 or Halo-Ku80 resulted in rapid loss of the other untagged Ku70 or Ku80 subunit, confirming that Ku70 and Ku80 are required to stabilize each other (Fig. 1B).

**Fig. 1.**
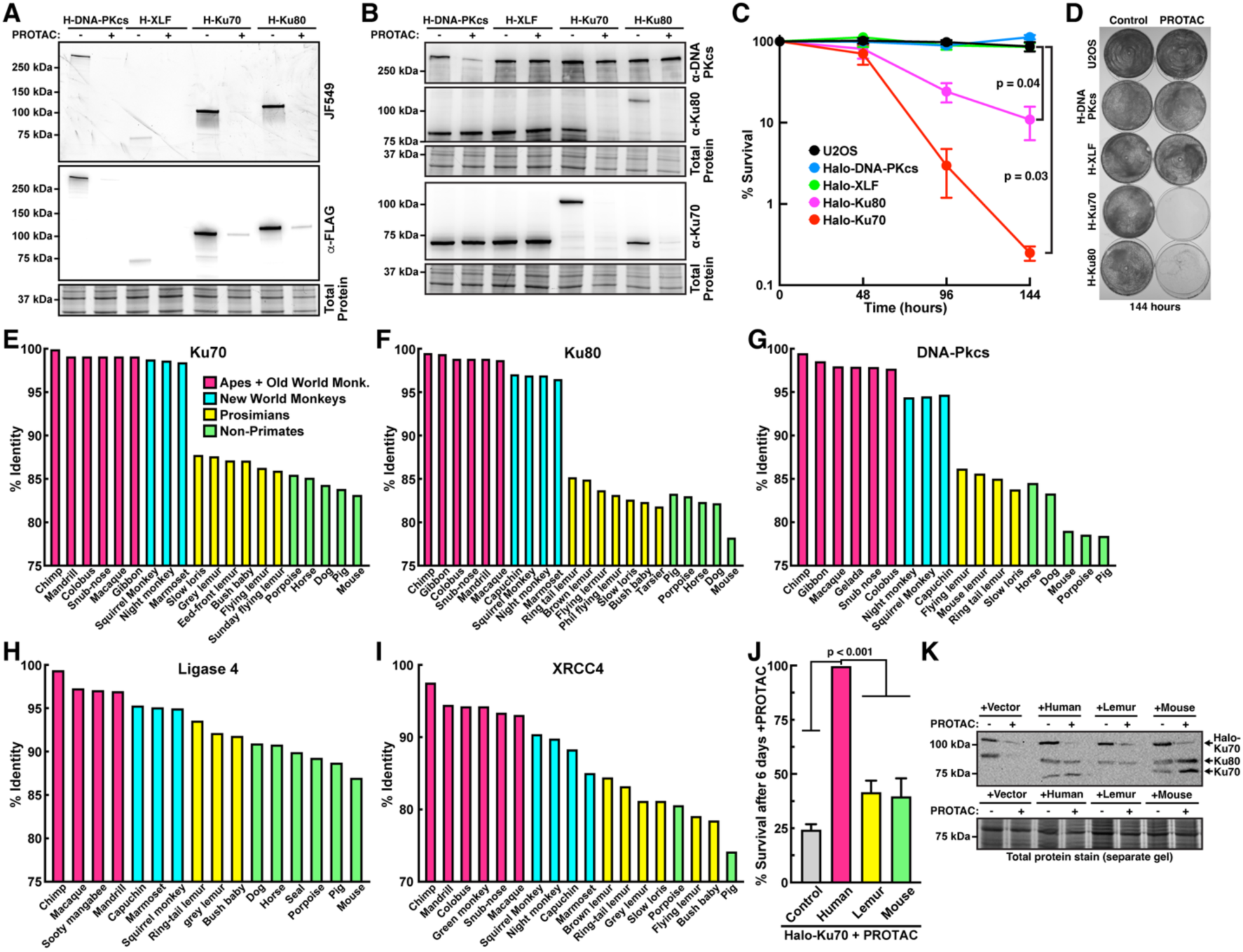
Cellular ablation of Ku70, or Ku80 but not XLF or DNA-PKcs results in rapid cell death in U2OS cells. **(A)** Fluorescence gel (top),anti-FLAG western blot (middle), and loading control (bottom) of cell lysates generated from U2OS cells expressing Halo-DNA-PKcs, XLF, Ku70, or Ku80 that had been incubated with 1 µM Halo-PROTAC or vehicle control for 48 hrs**. (B)** Western blots probed with antibodies against DNA-PKcs, Ku80, and Ku70 of cell lysates generated from U2OS cells expressing Halo-DNA-PKcs, XLF, Ku70, or Ku80 that had been incubated with 1 µM Halo-PROTAC or vehicle control for 72 hours. **(C)** Quantification of MTT cell viability assay of U2OS cells expressing Halo-DNA-PKcs, XLF, Ku70, or Ku80 that had been incubated with 1 µM Halo-PROTAC for the indicated time (N=3 biological replicates, Mean±SEM, T-test). **(D)** Cell viability analysis using crystal violet staining of U2OS cells expressing Halo-DNA-PKcs, XLF, Ku70, or Ku80 that had been incubated with 1 µM Halo-PROTAC for 6 days. **(E-I)** NHEJ factor orthologues of Ku70 (**E**), Ku80 (**F**), DNA-PKcs (**G**), Lig4 (**H**), or XRCC4 (**I**) from a variety of non-human primates (apes and old-world monkeys, red; new-world monkeys, blue; prosimians, yellow) and other mammals (dog, horse, seal porpoise, pig, and mouse; green) as indicated were compared to the human sequence. Percent identity for each to the human sequence is presented. **(J)** Rescue experiments for Halo-Ku70 expressing U2OS cells treated with 1 µM Halo-PROTAC for XX hours, complemented by stable transfection of lemur, mouse, or human Ku70. Cell survival was assessed by MTT staining and compared to blasticidin resistance conferred by the expression plasmids (N=3 biological replicates, Mean±SEM). **(K)** Western blots probing for Ku70 and Ku80 of Halo-Ku70 U2OS cells treated with 1 µM Halo-PROTAC, demonstrating that lemur and mouse Ku70 stabilize human Ku80 to confirm expression of Lemur Ku70 and complex formation with human Ku80.

Incubation of the five U2OS cell lines with 1 µM Halo-PROTAC resulted in significant loss of viability over the course of 6 days for cells expressing Halo-Ku70 or Halo-Ku80 but did not affect viability of parental cells or cells expressing Halo-XLF or Halo-DNA-PKcs (Fig. 1C,D). Halo-Ku70 cells died more rapidly than Halo-Ku80 cells (Fig. 1C, ∼3% of Halo-Ku70 or ∼24% of Halo-Ku80 are viable after 4 days, and ∼0.3% or ∼11% after 6 days). Although ∼70% of Halo-Ku70 cells were viable after 48 hours exposure to 1 uM Halo-PROTAC (Fig. 1C), cell viability was already impacted by a brief 6-hour exposure to the Halo-PROTAC, followed by replating in fresh media without PROTAC ligand (Fig. S1E), suggesting that even a temporary depletion of Ku results in severe changes in cellular processes from which cells cannot fully recover.

### Sequence comparisons suggest higher primate specific evolution of the three components of the DNA-PK complex, but not components of the ligase IV complex

To gain insight into possible functions of Ku in higher primates versus other mammals, we compared human NHEJ factor sequences to those of apes and old-world monkeys (red), new-world monkeys (blue), prosimians (yellow), and non-primates (porpoise, horse, dog, pig, mouse, green) (Fig. 1E-I). While sequence changes in XRCC4 and Ligase 4 appear to have evolved gradually, sequence identity in the subunits of DNA-PK abruptly changed (>10%) at the prosimian/new world monkey juncture (Fig. 1E-I). These substantial changes in DNA-PK polypeptide sequences, suggest novel properties of Ku70/80 and DNA-PKcs that may be specifically required in higher primates.

### Prosimian and Mouse Ku70 can only partially rescue Halo-Ku depletion in human cell lines

We next tested whether the higher primate homolog of Ku70 is required to support the cell-essential role of Ku70. To this end, a cDNA encoding Ku70 was isolated from a previously studied cell line from a ring-tailed lemur (prosimian) ^7^. Halo-Ku70 U2OS cells stably expressing Ku70 from either Human, Lemur, or Mouse using the piggy-bac system were tested for their ability to rescue cell viability after degradation of the endogenous human Halo-Ku70 (Fig. 1J). Human and Mouse Ku70 were expressed at similar levels to Halo-Ku70 and stabilized endogenous human Ku80 (Fig. 1K), confirming proper Ku heterodimer formation. Lemur Ku70 also stabilized human Ku80 to a similar degree as the Human and Mouse Ku70 transgenes, confirming that it is expressed and can associate with human Ku80 (Fig. 1K). However, the Ku70 antibody was unable to recognize the Lemur Ku70 protein, which did not allow us to determine its precise expression levels. Neither Lemur nor Mouse Ku70 expression rescued cell viability after depletion of human Halo-Ku70 (Fig. 1J). To further confirm this observation, we inserted the Flag-HaloTag at the N-terminus of Ku70 in HEK293T cells (Fig. S1F). Similar to U2OS cells, treatment of HEK293T cells expressing Halo-Ku70 with the Halo-PROTAC ligand significantly reduced cell viability (Fig. S1G,H), which was not fully rescued by expression of Lemur or Mouse Ku70 (Fig. S1I,J). In addition, the HEK293T Halo-Ku70 cells also allowed us to assess the ability of the Ku complex to support NHEJ in VDJ recombination assays. Halo-Ku70 degradation using the Halo-PROTAC ligand significantly reduced NHEJ activity in HEK293T cells (Fig. S1K,L). Expression of Human, Mouse, or Lemur Ku70 fully restored NHEJ activity, while Human Ku70 with an N-terminal deletion, known to have an NHEJ defect, failed to rescue VDJ recombination of depletion of the endogenous Halo-Ku70 (Fig. S1K,L). We conclude that Human Ku70 has specific functional features in higher primates, which are not present in prosimian or rodent Ku70, that are important for Ku’s essential role in human cells. Furthermore, our results show that Lemur and Mouse Ku70 are fully functional to support NHEJ in HEK293T cells, demonstrating that reduced NHEJ activity does not underlie the loss of viability after Ku70 depletion in human cells.

### Gene expression is significantly altered after 48 hours of Ku70 degradation in U2OS cells, but no activation of an innate immune response is observed

Zhu et al. reported that Ku80 depletion in HCT116 cells dramatically upregulates innate immune signaling via the STAT1 and protein kinase R (PKR) pathways ^15^. They proposed that Ku functions to restrain abundant Alu RNA molecules by inhibiting their ability to bind innate immune sensors that recognize viral double-stranded RNA. To investigate the impact of Ku degradation on gene expression in U2OS cells, we carried out RNA-seq experiments comparing control Halo-Ku70 or Halo-DNA-PKcs cells to cultures treated with 1 µM Halo-PROTAC for 48 hours, the timepoint immediately prior to rapid cell death of Ku70 depleted cells. No significant changes in gene expression were observed after Halo-DNA-PKcs degradation (Fig. S2A). In contrast, transcription was significantly altered in U2OS Halo-Ku70 cells treated with 1 µM Halo-PROTAC compared to controls (Fig. 2A).

**Fig. 2.**
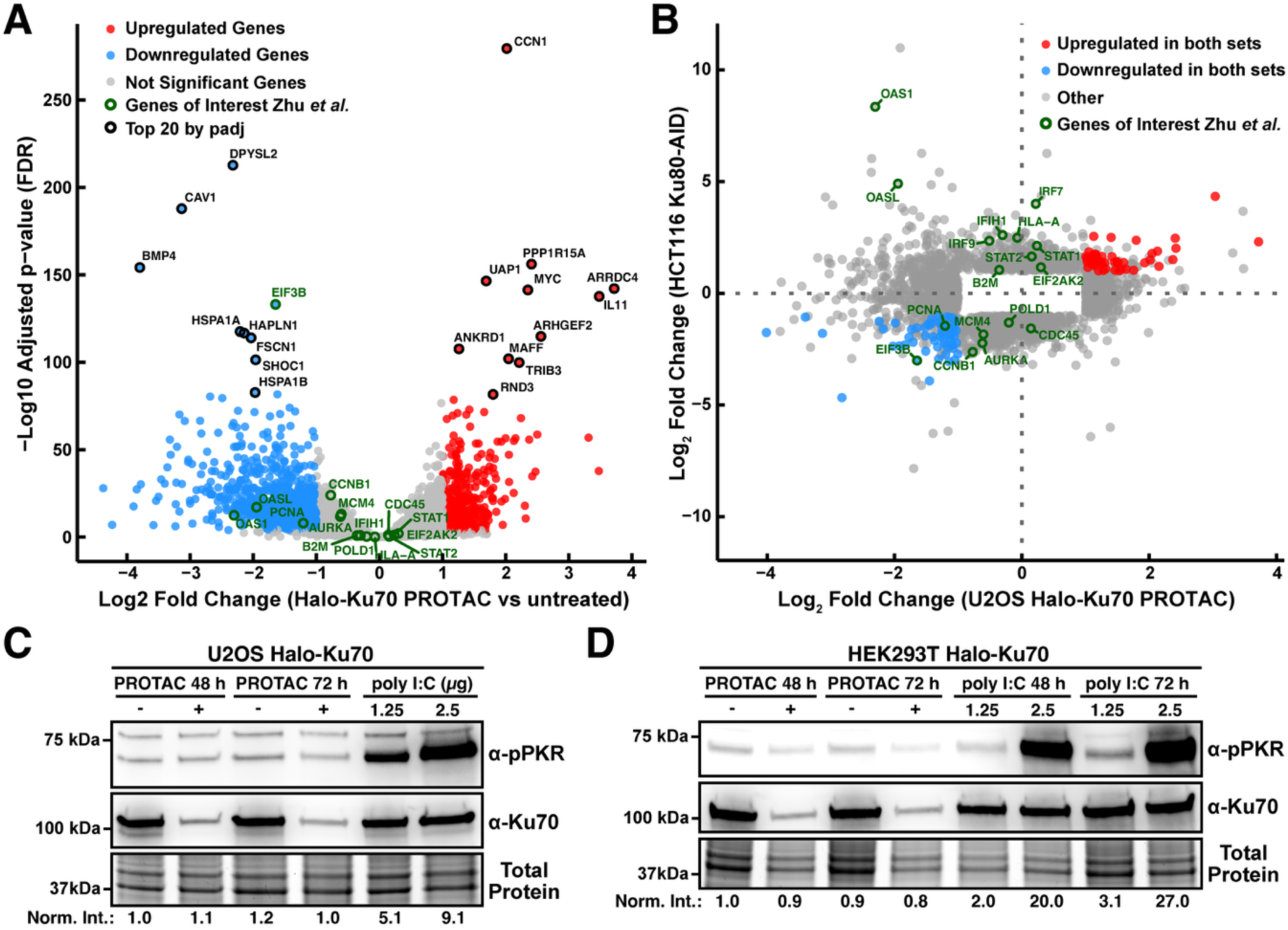
Gene expression is markedly altered within 48 hours of Ku70 ablation, but no obvious activation of innate immune pathways is observed. (A) Volcano plot comparing mRNA levels in untreated Halo-Ku70 cells to cells treated with 1 uM Halo-PROTAC for 48 hours. Genes highlighted in green denote genes observed to be significantly altered after Ku80 degradation in HCT116 cells in previous work by *Zhu et al.*. (B) Scatter plot of the Log2 fold changes of mRNA levels in U2OS Halo-Ku70 cells and HCT116 Ku80-AID cells after Ku depletion relative to control cells. Genes highlighted in green denote genes observed to be significantly altered after Ku80 degradation in HCT116 cells in previous work by *Zhu et al.*. (C-D) Immunoblot demonstrating induction of T446 phosphorylation on PKR by transfection of poly I/C but not by 1 uM Halo-PROTAC treatment of (C) Halo-Ku70 U2OS cells or (D) Halo-Ku70 HEK293T cells. Intensity of pPKR signal was normalized to total protein loading.

We hypothesized that loss or gain of specific transcripts after Ku ablation in diverse cell types might reveal important mechanistic insight into the basis of Ku’s essential nature in human cells. After Ku70 degradation in U2OS cells 527 genes were significantly upregulated and 907 genes were significantly downregulated (Fig. 2A, Fig. S2B,C). When comparing our result to previously published RNAseq data by Zhu *et al*. after Ku80 degradation in HCT116, no strong correlation between genes with altered RNA levels was detected between the two cell types (Fig. 2B). Importantly, we did not observe the upregulation of STAT1 dependent innate immunity genes in U2OS cells, which was observed in HCT116 cells (e.g. EIF2AK2, Fig. 2A,B). In contrast, key factors involved in double-stranded RNA sensing (e.g. OASL, OAS1), which were found to be upregulated in HCT116 cells after Ku80 depletion, were significantly downregulated in U2OS cells after Halo-Ku70 degradation (Fig. 2B). To further analyze the activation of double-stranded RNA sensing in U2OS and HEK293T cells after Halo-Ku70 degradation, we assessed phosphorylation of PKR (EIF2AK2) at T446. Transfection of Halo-Ku70 expressing U2OS and HEK293T cells with double-stranded RNA (polyI:C) but not Halo-Ku70 degradation led to robust induction of PKR phosphorylation (Fig. 2C,D), demonstrating that this double-stranded RNA sensor is not activated in U2OS and HEK293T cells lacking Ku70. We conclude that activation of innate immune signaling pathways by cytoplasmic double-stranded RNA is unlikely to contribute to cell death in U2OS or HEK293T cells after degradation of Ku70.

The essential translation initiation factor EIF3B was significantly downregulated in HCT116 and U2OS cells after Ku subunit depletion (Fig. 2A,B, Fig. S2D). EIF3B protein levels were also reduced in Ku70 depleted HEK293 cells as assessed by mass-spectrometric methods by Kelly et al.^14^. DepMap co-dependency analysis of RNAi experiments revealed that EIF3B, Ku70, and Ku80 are highly correlated (Fig. S2D,E), which would be expected if the loss of EIF3B expression contributes to the reduction in cell viability after Ku degradation.

### Ablation of either Ku subunit results in dysregulation of RNA splicing

Since we did not observe activation of innate immune signaling in U2OS cell after Ku70 depletion, we considered alternative explanations for the rapid cell death observed. Our RT-PCR validation experiments yielded unexpected products in Ku-ablated cells, potentially resulting from alterations in splicing. Alu elements have been widely implicated in mRNA splicing ^23–28^, and because Ku was recently demonstrated to specifically bind antisense Alu RNAs ^15,29^, we analyzed alternative splicing after Ku70 ablation in U2OS cells using LeafCutter ^30^. LeafCutter quantifies alternative splicing directly from RNA-seq split-read counts by overlapping introns into clusters of competing splicing events and testing whether intron usage differs between two conditions. For each intron, LeafCutter reports two effect-size metrics: |ΔPSI|, the absolute change in percent-spliced-in (the fraction of the cluster’s reads attributed to that intron) between conditions, and logef, the log-scale effect size from LeafCutter’s Dirichlet-multinomial regression model, which is proportional to the log-odds of intron usage and is more sensitive to changes in low-abundance introns.

Degradation of Halo-Ku70 in U2OS cells results in significant changes in mRNA splicing (Fig. 3A), with approximately 9% of expressed multi-exon genes having at least one alternatively spliced intron cluster (Fig. S3A). This impact of Ku depletion on U2OS transcriptome is comparable to that of well-characterized ubiquitous alternative splicing regulators^18,31,32^. To confirm these findings, we analyzed alternative splicing in RNA-seq data sets after Ku80 depletion in HCT116 and Ku70 depletion in HEK293 cells published by Zhu. *et al.* ^15^. This analysis demonstrated that splicing was also significantly altered in both HCT116 and HEK293 cells after Ku80 or Ku70 degradation (Fig. 3B,C), impacting 4.6% and 12.3% of expressed multi-exon genes (Fig. S3A), respectively. A classification of splicing event types (“SE”, cassette exon; “A5SS”, alternative 5’ splice site; “A3SS”, alternative 3’ splice site; “Other” for other types) demonstrated that cassette exon was the most frequent splicing alteration in all datasets (>75% of splicing changes for all cell lines, Fig. 3D), impacting 7.8%, 3.6%, and 11% of expressed genes in U2OS, HCT116, and HEK293 cells after Ku70 or Ku80 depletion (Fig. S3B), respectively. Cassette exons potentially repressed by Ku, and therefore more spliced into transcripts after Ku depletion (“Ku-repressed”), were more abundant in U2OS and HCT116, while cassette exons that were spliced out more frequently when Ku was degraded from cells (“Ku-activated”) were predominant in HEK293 (Fig. 3E). 72 genes, including several essential genes, with at least one alternatively spliced intron cluster were shared across the three cell lines (p=5.85e-44, 3-way intersection, Superexact test), suggesting a common mechanism underlying these changes (Fig. 3F, Table S1). GO enrichment analysis across all three cell lines revealed that Ku depletion preferentially affected splicing of mRNAs encoding proteins involved in a variety of cell essential processes (e.g. nuclear transport, ribosome biogenesis, splicing), with ribosome biogenesis factors being strongly enriched in the genes with altered splicing in all three cell lines after Ku ablation (Fig. 3G-J). Since genes contributing to RNA processing and splicing were enriched in alternatively spliced RNAs in all three cell lines, it is possible that depletion of splicing factors exacerbates the dysregulated splicing that occurs in the absence of Ku. Thus, some observed changes may not be a direct consequence of Ku depletion, per se, but are an indirect consequence of altered protein levels of splicing factors.

**Fig. 3.**
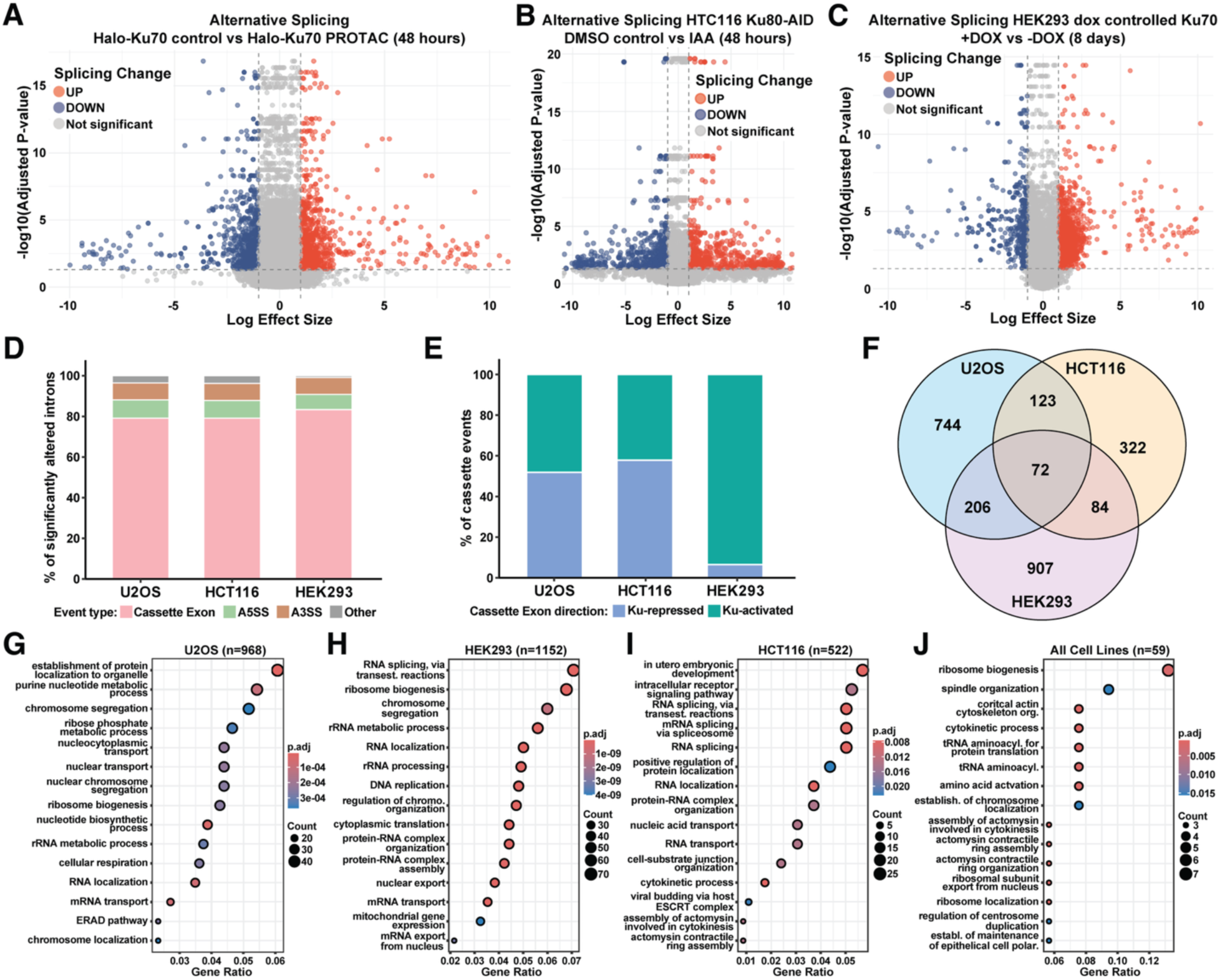
Ablation of either Ku subunit results in dramatic dysregulation of RNA splicing. (A-C) Volcano plots showing differential splicing events identified by LeafCutter in (A) Ku70-depleted U2OS cells, (B) Ku80-depleted (by IAA addition) HCT116 cells (*Zhu. et al.*), and (C) Ku70-depleted (by doxycycline withdrawal) HEK293 cells (*Zhu et al.*). Each point represents an intron cluster, with significantly up-and downregulated events (|logef| > 1, adjusted P < 0.05) shown in red and blue, respectively; non-significant events are in grey. Dashed lines indicate effect size and significance thresholds. **(D)** Classification of Ku-dependent alternative splicing event classes in U2OS, HCT116 and HEK293 (**E)** Relative proportion of directions for the “cassette exon” event class; “Ku-repressed”: cassette exons more spliced-in after Ku depletion; “Ku-activated”: cassette exons more spliced-out after Ku depletion. **(F)** Venn diagram showing the overlap of genes with altered splicing after Ku depletion in U2OS, HCT116, and HEK293 cells. **(G-J)** Gene ontology analysis of biological processes for genes with alternatively spliced introns in (G) U2OS cells 48 hours after Halo-Ku70 depletion, (H) HEK293 cells with doxycycline controlled Ku70 expression 8 days after doxycycline withdrawal, (I) HCT116 Ku80-AID cells 48 hours after IAA addition, and (J) introns alternatively spliced in all three cell lines.

In total, these results demonstrate that splicing is significantly altered in three independent cell lines after Ku depletion using distinct methodologies. The number of genes with altered splicing after Ku depletion is comparable to the loss of ubiquitous alternative splicing regulators and cassette exons are the most common splicing change observed after Ku depletion. We therefore propose that the Ku complex is a bona-fide regulator of mRNA splicing, predominantly controlling splicing of cassette exons.

### The Ku complex regulates splicing of antisense Alu element containing introns

Since Ku selectively binds Alu elements, with a strong bias for repeats in antisense orientation ^15^, and because Alu elements have been widely implicated in mRNA splicing and evolution of new exons ^33,34^, we hypothesized that cryptic splice sites within Alu sequences may be more accessible upon Ku depletion. Consistent with this hypothesis, splice junctions of Ku-repressed cassette exons overlapped with Alu elements at significantly elevated rate compared to non-significant introns in all three cell lines (Fig. 4A, S3C,D; Fisher’s exact test, OR=1.8–5.4, BH-adjusted p≤1.3×10⁻⁵). Furthermore, Alu elements that overlap with alternative splice junctions of cassette exons were almost exclusively (94-100%) found in the antisense orientation to the host gene (Fig. 4A). Junctions of Ku-activated exons did not significantly overlap with Alu elements (Fig. 4A), suggesting a different mechanism for their exclusion in absence of Ku. One potential explanation would be looping out of the exon by base-pairing of inverted Alu pairs (irAlu) flanking the excised exons ^35^. Indeed, Ku-activated cassette exons are significantly enriched for exon-spanning irAlu pairs in all three cell lines (Fisher’s exact test OR=2.0-3.7, p≤3.5×10⁻⁵; Fig. 4B), suggesting that Ku binding to the asAlu element within the irAlu pair may prevent base-pairing of exon spanning irAlu elements to promote inclusion of these exons.

**Fig. 4.**
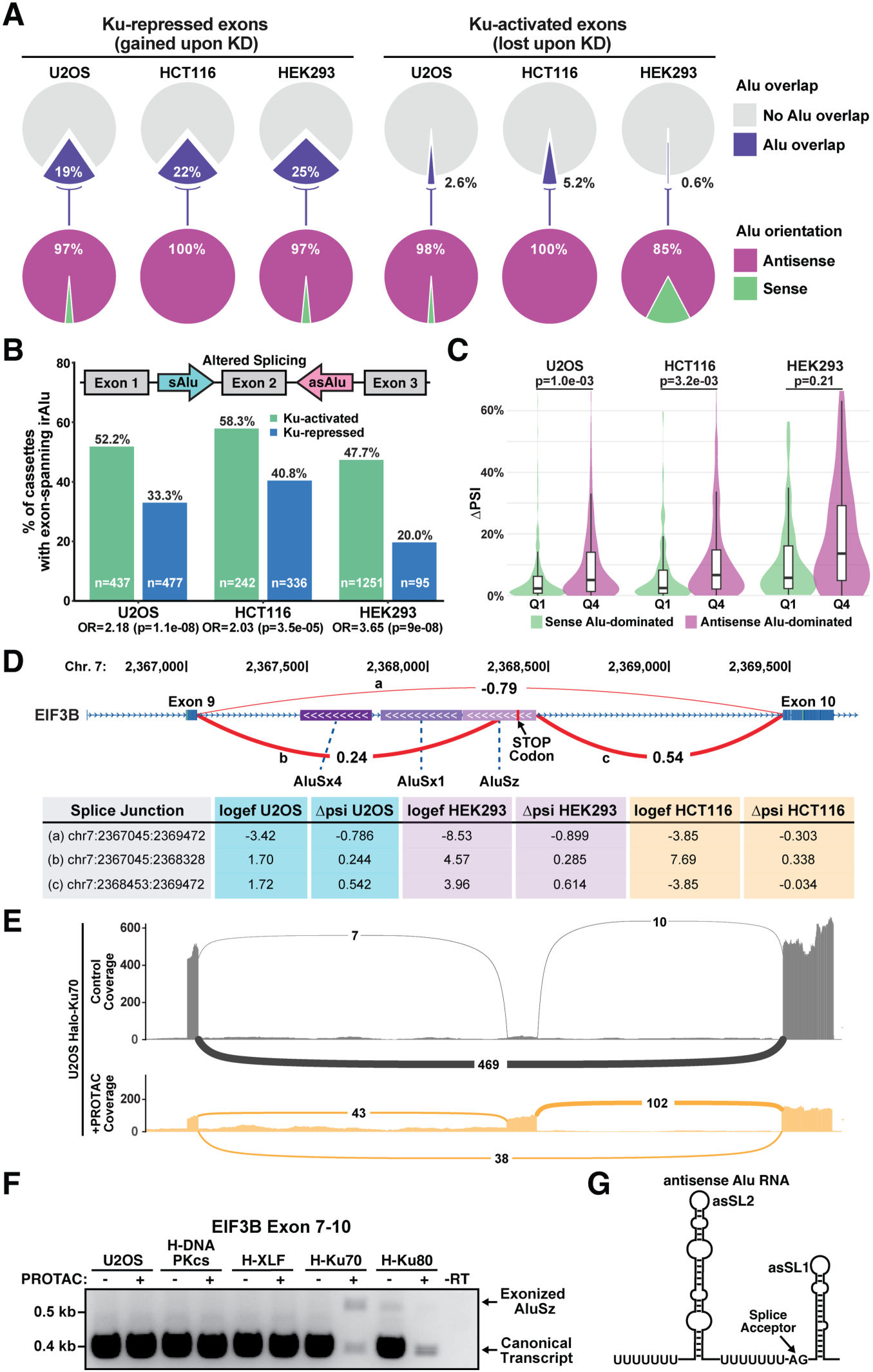
Ku depletion activates cryptic splice sites within antisense Alu elements. **(A)** Relative proportion (top) of splicing junctions of Ku-repressed and Ku-activated cassette exons overlapping intronic Alu and (bottom) orientation of intronic Alu overlapping splice junctions of Ku-repressed and Ku-activated cassette exons, relative to host gene. **(B)** Quantification of the fraction of Ku-activated and Ku-repressed cassette exons flanked by inverted Alu repeats (irAlu). Over representation of exon-spanning irAlu in Ku-activated versus Ku-repressed cassettes was tested per dataset by two-sided Fisher’s exact test on the 2×2 (architecture × direction) contingency table. **(C)** ΔPSI of Ku-repressed cassette exons stratified by per-intron antisense/sense Alu ratio (±2 kb around splice junctions). Q1=sense-dominated quartile, Q4=antisense-dominated quartile; p= Wilcoxon rank-sum, BH-adjusted across datasets. **(D)** Leafcutter analysis of splicing between exon 9 and 10 of the *EIF3B* gene in Halo-Ku70 U2OS cells treated with Halo-PROTAC relative to control cells. Letters (a,b,c) denote splice junctions connected by the arcs indicating ΔPSI after Ku depletion. Purple shaded boxes depict the position of antisense Alu elements (white arrows indicate orientation). Table (bottom) contains Leafcutter outputs for all three cell lines after Ku depletion. **(E)** Sashimi plot of RNA-seq read coverage and splice junction usage in at the *EIF3B* locus in control (top, grey) and Halo-PROTAC treated Halo-Ku70 U2OS cells (bottom, orange). Numbers and arc thickness reflect junction read counts. **(F)** RT-PCR analysis of EIF3B mRNA splicing from exon 9 to 10 after depletion of various NHEJ factors in U2OS cells. **(G)** Model and sequence of antisense Alu RNAs.

Since Ku has a strong binding preference for antisense Alu elements, we tested whether the orientation of nearby Alu elements predicts the magnitude of the splicing change (measured by |ΔPSI|). Ranking of Ku-repressed cassettes in quartiles based on their antisense/sense Alu content ratio shows |ΔPSI| scales quantitatively with the antisense Alu content, and the most antisense-enriched quartile undergoes 2.2-to 2.7-fold larger absolute PSI changes than the most sense-dominated quartile in all three cell lines (Fig. 4C). To further confirm these findings, we carried out RT-PCR analysis across multiple exon junctions for the *UPF3*, and *MELK* genes (Fig. S3E-H). This RT-PCR analysis also demonstrated altered splicing, including skipping, of exons flanking introns containing antisense Alu elements in Halo-Ku70 depleted U2OS cells.

Together, these results demonstrate that Ku depletion results in splicing alterations in sequences containing antisense Alu elements and altered splicing junctions frequently overlap directly with antisense Alu elements.

### Ku suppresses the use of cryptic splice sites within antisense Alu elements

To define the molecular nature of Ku-sensitive splicing events, we focused on cryptic splice junctions whose usage increased upon Ku70 depletion. The essential *EIF3B* gene, for which RNA levels were downregulated in all three cell lines, contains two novel splice junctions between exons 9 and 10, which were upregulated after Ku70 depletion at the expense of the canonical junction that properly connects these exons (Fig. 4D,E). This alternative splicing event is a canonical example of a Ku-repressed cassette exon type, with the splice acceptor and donor sites residing within an antisense Alu element in the intron intervening exons 9 and 10 of the *EIF3B* gene (Fig. 4D). Precisely the same altered splice junctions within the Alu element were also detected in HCT116 and HEK293 cells after Ku depletion (Fig. 4D, Fig. S3I). We confirmed the alternative splicing event by RT-PCR (Fig. 4F). Antisense Alu RNAs contain two stem loops (asSL2 and asSL1) preceded by poly-pyrimidine stretches (Fig. 4G). Poly-pyrimidine stretches followed by an AG dinucleotide are the key sequence features of splice acceptor sites (Fig. 4G), and the AG dinucleotide preceding asSL1 within the antisense AluSz element in intron 9 of the *EIF3B* gene is used as the splice acceptor in the cassette exon upregulated after Ku depletion (Fig. 4D,E,G).

A small amount of the transcript containing the cassette exon within the antisense Alu element in intron 9 of the *EIF3B* gene was also detected by RT-PCR in control Halo-Ku80 cells (Fig. 4F); similar splicing alterations were observed in RT-PCR analyses of other transcripts in Halo-Ku80 cells. We suggest that the HaloTag on the N-terminus of Ku80 impacts its ability to limit aberrant splicing, providing a likely explanation for the fitness deficits in cells expressing Halo-Ku80. All other splice junctions within the EIF3B transcript were unchanged after Ku depletion (Fig. S3I,J) in U2OS, HCT116, and HEK293 cells, even though antisense Alu elements are present in several other introns of the gene (Fig. S3I). This demonstrates that the altered splicing caused by Ku depletion is specific to certain introns containing antisense Alu elements, rather than a general defect in mRNA processing. Importantly, we identified additional examples of cassette exons within antisense Alu elements which were derepressed by Ku depletion in all three cell lines (Fig. S4A-E), suggesting that the inclusion of cassette exons within antisense Alu elements is a general mechanism underlying the splicing alterations observed after Ku ablation. In some examples, the cryptic splice acceptor sites (whose use increased after Ku depletion) were upstream of asSL2 in antisense Alu elements (Fig. S4B,D,E); these are also preceded by poly-uridine stretches (Fig. 4G) and shown to associate with Ku in CLIP-Seq experiments ^15^. Finally, we identified a gene (*NAA35*) containing an alternative 3’-splice sites within an antisense Alu element upregulated after Ku depletion (Fig. S4F), and examples of exons flanked by irAlu elements which were more frequently skipped after Ku ablation[confirmed by RT-PCR (Fig. S4G-J)].

To infer the possible consequences of mis-splicing caused by Ku ablation, we tested how frequently the cryptic inclusion of Ku-repressed cassette exons introduces a premature termination codon (PTC) into the coding sequence of protein-coding genes, making the resulting transcripts potential targets of nonsense-mediated decay (NMD). Of 908 Ku-repressed cassettes analyzed across the three datasets, 530 (54-82% per dataset) fell within the coding sequence (CDS) of an exon-skipping transcript isoform; 134 fell in an untranslated region (UTR) and 244 could not be assigned to such an isoform (Fig. S4K). Among in-CDS cassettes, 82-91% were predicted to trigger NMD under the 50-nt NMD rule, corresponding to 447 of 908 Ku-repressed cassettes and 301 unique genes, including *EIF3B* (Fig. S4L). Cryptic inclusion of Ku-repressed cassettes is therefore predicted to generate NMD-substrate transcripts for approximately half of all such events, consistent with the reduced RNA levels observed for EIF3B mRNA.

Altogether, these observations demonstrate that Ku depletion activates cryptic splice sites within intronic antisense Alu elements, and ∼50% of the resulting cassettes introduce a premature termination codon predicted to elicit nonsense-mediated decay.

### Mis-splicing of the EIF3B mRNA contributes to cell death after Ku70 depletion

Ku depletion leads to mis-splicing and reduced abundance of the EIF3B mRNA (Fig. 2 and Fig. 4). Importantly, inclusion of the cassette exon within the antisense Alu element contained in intron 9 of the *EIF3B* gene introduces a premature stop codon in the coding sequence of the mRNA (Fig. 4D), and indeed this miss-spliced EIF3B isoform was identified as a likely target of NMD by our stringent NMD prediction analysis.

To confirm that the antisense AluSz element in intron 9 of the *EIF3B* gene containing the cassette exon upregulated after Ku70 degradation (Fig. 4D) is necessary for mis-splicing and RNA degradation after Ku70 depletion, we deleted this sequence from at least one of the *EIF3B* alleles using Cas9 and two guide RNAs targeting intron 9 of the EIF3B gene in U2OS cells (Halo-Ku70 ΔAlu, Fig. 5A,B). Removal of the AluSz element from intron 9 of the *EIF3B* gene did not impact expression of the mature EIF3B mRNA in Halo-Ku70 U2OS cells under control conditions relative to parental U2OS cells (Fig. 5C). In contrast, the EIF3B mRNA level was reduced ∼200-fold (0.5% of parental U2OS cells) in Halo-Ku70 treated with Halo-PROTAC ligand and removal of the AluSz element from intron 9 of the *EIF3B* gene increased EIF3B expression ∼15-fold to 7.5% of the RNA levels detected in parental U2OS cells (Fig. 5D). This modest rescue of the EIF3B mRNA levels likely reflects our inability to remove the AluSz element from intron 9 of all *EIF3B* alleles. Consistent with this interpretation, analysis of the expression level of the EIF3B transcript containing the cassette exon spanning the antisense AluSz element was significantly increased to a similar degree in Halo-Ku70 and Halo-Ku70 ΔAlu U2OS cells (Fig. 5E). Importantly, removal of the AluSz element from intron 9 of at least one of the EIF3B alleles was sufficient to significantly increase cell survival 96 hours after Ku70 depletion compared to Halo-Ku70 U2OS cells in three independent clones (Fig. 5,F). Extended cultures of the Halo-Ku70 U2OS ΔAlu cells in the presence of the Halo-PROTAC ligand revealed that the cells stop growing by day 9 demonstrating that the partial rescue of EIF3B mRNA expression in Ku depleted cells is not sufficient to supportcell viability. This suggests that that loss of EIF3B is in part (but not fully) responsible for the reduction of human cell viability in the absence of Ku.

**Fig. 5.**
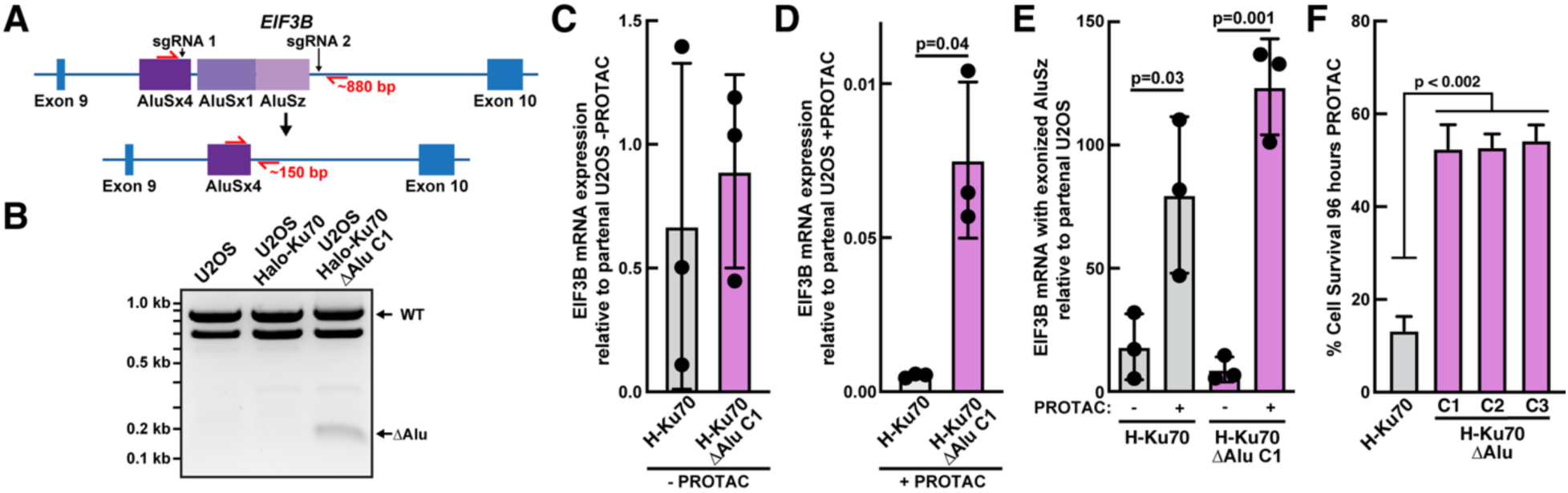
Antisense Alu RNA directly associates with Ku70/80 and its exonization destabilizes the EIF3B mRNA. **(A)** Experimental approach to delete the antisense AluSz element from intron 9 of the *EIF3B* gene. **(B)** Agarose gel of genomic PCR products from U2OS, Halo-Ku70 U2OS, and Halo-Ku70 ΔAlu U2OS cells using the primers indicated in (A). **(C-E)** Quantification of RT-qPCR of RNA extracted from parental U2OS, Halo-Ku70 U2OS and Halo-Ku70 ΔAlu U2OS cells (C) under control conditions and (D) after PROTAC treatment using a primer specific to the mature EIF3B mRNA, or (E) using a primer set specific to the splice junction generated by AluSz exonication in control and PROTAC treated cells. All Ct values were normalized to GAPDH and Halo-Ku70 U2OS and Halo-Ku70 ΔAlu U2OS RNA levels were normalized to that of parental U2OS cells (N=3 biological replicates, Mean±SD, T-Test). **(F)** Quantification of cell viability of U2OS cells expressing Halo-Ku70 and Halo-Ku70 ΔAlu U2OS cells (N=3 biological replicates, Mean±SEM, T-Test).

### Ku stabilizes asSL1 stem loop in cells

Previous results by other have shown that the Ku complex preferentially associated with asSL1 of antisense Alu elements in cells using irCLIP experiments^15^. In addition, they demonstrated that Ku directly associates with asSL1 *in vitro*, however this binding does not explain the specificity for antisense Alu RNA in cells, since Ku bound to the sense SL1 stem loop with similar affinity ^15^. Ku binding to asSL1 in antisense Alu RNAs could physically block association of the splicing machinery with the splice acceptor site immediately upstream. Alternatively, the folding of asSL1 into a stem loop could be sufficient to prevent recognition of the splice acceptor site and Ku binding could promote the formation of or stabilize the asSL1 stem loop. To determine whether Ku regulates asSL1 stem loop folding in cells we carried out dimethyl sulfate (DMS) mutational profiling (MaP) of the antisense AluSz element in intron 9 of the *EIF3B* gene ^36^. DMS specifically reacts with unpaired Cytosine and Adenine nucleotides in cellular RNAs. DMS adducts are then converted into mutations in a cDNA sequence via relaxed fidelity reverse transcription which is subsequently read out by high-throughput sequencing (Fig. 6A). A high mutational frequency in DMS-MaP experiments identifies nucleotides located in single-stranded RNA regions, or bulges and loops within stem loop structures. In contrast, low DMS reactivity is consistent with nucleotides being engaged in base-pairing for instance in the context of stems within stem loops.

**Fig. 6.**
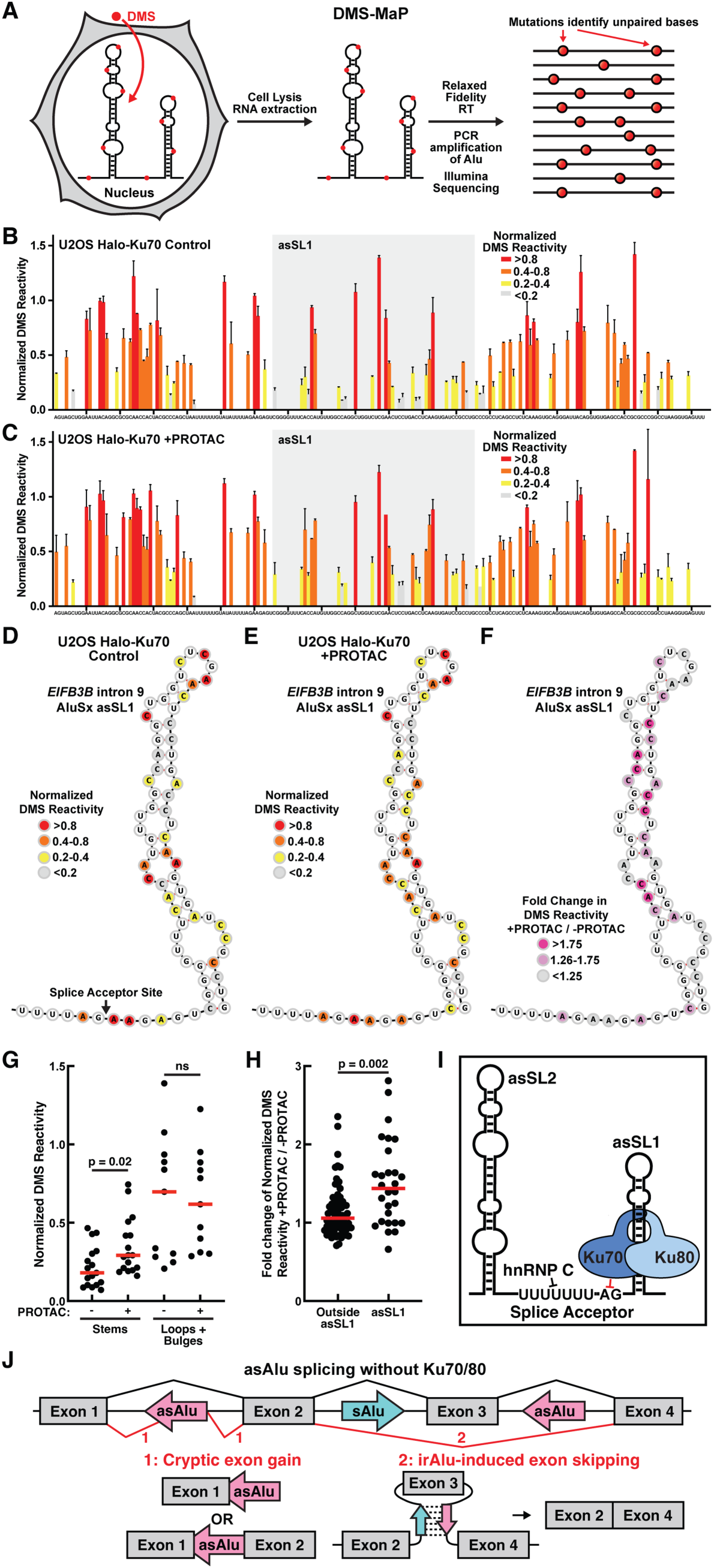
The Ku complex stabilizes the asSL1 stem loop within an antisense Alu element RNA. **(A)** Experimental approach for mapping RNA secondary structure in cells using DMS-MaP. **(B-C)** Normalized DMS-MaP reactivity profiles for an RNA amplicon from the antisense AluSz element in intron 9 of the *EIF3B* gene spanning the asSL1 stem loop from **(B)** control and **(C)** Halo-PROTAC treated (48 hours) Halo-Ku70 U2OS cells. Reactivity was normalized to the mean of the top 10% mutational frequencies observed in each replicate. Values plotted are the mean ± SD of two experimental replicates. Grey box indicates the asSL1 sequence. **(D-E)** Normalized DMS reactivity from **(D)** control and **(E)** Halo-PROTAC treated (48 hours) Halo-Ku70 U2OS cells mapped onto the secondary structure of the asSL1 stem loop. **(F)** Fold change of normalized DMS reactivity of Halo-PROTAC treated Halo-Ku70 U2OS cells relative to untreated controls mapped onto the secondary structure of the asSL1 stem loop. **(G)** Comparison of normalized DMS reactivity in paired stems and unpaired loops and bulges between control and Halo-PROTAC treated Halo-Ku70 U2OS cells. Each data point corresponds to an individual nucleotide, and the red bar shows the mean reactivity. Statistical significance was assessed by t-test. **(H)** Comparison of the fold change of normalized DMS reactivity of Halo-PROTAC treated Halo-Ku70 U2OS cells relative to untreated controls within asSL1 and in outside of asSL1. Each data point corresponds to an individual nucleotide, and the red bar shows the mean reactivity. Statistical significance was assessed by t-test. **(I)** Model of the structure of an antisense Alu element bound by Ku70/80 to block the cryptic splice acceptor site**. (J)** Model depicting potential outcomes of Alu element exonization or exon skipping due to the use of cryptic splice acceptors within antisense Alu elements de-repressed or the presence of irAlu elements after Ku depletion.

To amplify the antisense AluSz element in intron 9 of the *EIF3B* gene we carried out the reverse transcription step with a specific primer within intron 9 downstream of the AluSz element and then PCR amplified an amplicon covering 185 nucleotides of the AluSz element spanning asSL1. DMS reactivities from individual experimental replicates correlated well, demonstrating the reproducibility of the methodology (Fig. S5A,B). The overall DMS reactivity profile in control U2OS Halo-Ku70 cells was consistent with the canonical asSL1 stem loop structure (Fig. 6B,D), which was previously shown to bind to the Ku complex *in vitro* ^15^. Ku degradation led to a significant increase in the DMS reactivity of nucleotides within base-paired stems in asSL1, while the reactivity of unpaired nucleotides was unchanged (Fig. 6B-G). This observation demonstrates that the asSL1 stem loop unfolds more frequently in the absence of the Ku complex, suggesting that Ku binding to asSL1 stabilizes its folded state. The alterations in DMS reactivity were confined to the asSL1 sequence (62 nucleotides, mean fold change in DMS reactivity = 1.5), while the reactivity of the remainder of the antisense AluSz element sequence contained in the amplicon analyzed were largely unchanged (123 nucleotides, mean fold change in DMS reactivity = 1.1) (Fig. 6H, Fig. S5C). This demonstrates that Ku degradation specifically alters the secondary structure of the asSL1 stem loop rather than leading to general folding changes in the antisense Alu RNA.

In total, our DMS-MaP experiments demonstrate that Ku degradation results in increased unfolding of asSL1 within the antisense AluSz element in intron 9 of the *EIF3B* gene, which is consistent with a direct interaction of the Ku complex with asSL1 in cells. In addition, these results suggest that unfolding of asSL1 could allow the splicing machinery to access the cryptic splice acceptor preceding the RNA stem loop, providing a mechanistic explanation for the increased Alu element exonization in the absence of the Ku complex.

## DISCUSSION

Alu invasion of primate genomes is a double-edged sword. It is well-appreciated that NAHR promoted by Alu elements functioned to diversify primate genomes allowing for the evolution of traits like full color vision, advanced neuronal development and function (reviewed in ^37^, lack of tails in apes ^38^, etc.). It is similarly well-documented that detrimental Alu-mediated recombination events are the basis of numerous genetic diseases (for example hemophilia, familial hypercholesterolemia, spinal muscular atrophy, etc.) ^39,40^. Moreover, Alu-mediated somatic recombination events promote many types of human cancers (in genes like BRCA1, BRCA2, MSH2, MLL1) (reviewed in ^10,41^). On the other hand, the presence of Alu elements in transcribed mRNAs further diversifies the potential transcriptome by providing a robust mechanism to alter protein products by mRNA splice variation, as has been appreciated for several decades ^23–28^.

It appears that the acquired ability to diversify protein expression by insertion of Alu elements in protein coding genes is apparently too robust – necessitating co-evolution of a mechanism to regulate alterations in splicing. The expansion of Alu elements in primate genomes resulted in large increases in the cellular levels of the SRP Alu-binding heterodimer SRP9/14 as well as components of the DNA-PK complex ^7^. In addition, hnRNP C and DHX9 have been shown to inhibit intronic Alu elements from altering host gene splicing ^18,20^. Our data presented here demonstrate that Ku depletion leads to large scale changes in splicing, comparable in magnitude to loss of abundant alternative splicing regulators ^18,31,32^. We observed both exonization of antisense Alu elements and skipping of exons flanked by inverted Alu element pairs (irAlus). The simplest model to explain how Ku regulates splicing, is that Ku (with its relatively weak dsRNA binding activity) binds intronic antisense Alu elements in immature RNA transcripts, modulating potential alternative splice sites or secondary structure formation by irAlus. Consistent with this model, Ku has been demonstrated previously to associate with both RNA, RNA processing factors ^42–44^ and with the spliceosome ^45,46^.

Ku was previously shown to predominantly bind to the second stem loop in antisense Alu RNA in cells (asSL1, Fig. 6I) ^15^. Importantly, asSL1 is immediately preceded by the cryptic splice acceptor site which is de-repressed in exonized antisense Alu elements after Ku depletion (Fig. 6I). Splice acceptors contain two critical sequence elements, a poly-pyrimidine tract followed by the invariant AG dinucleotide, which marks the end of the intron ^47^. Both of these elements are present in antisense but not sense Alu elements. The poly-pyrimidine tract is bound by the splicing factor U2AF2 ^47^, and elegant work has demonstrated that hnRNP C competes with U2AF2 to prevent the usage of cryptic splice acceptors within antisense Alu elements ^18^. The AG dinucleotide is recognized by U2AF1, which also associates with the first two nucleotides of the subsequent exon ^48^. In cryptic splice acceptors that are upregulated after Ku degradation the AG site is located within a few nucleotides of the Ku70/80 binding stem loops. The stem loop structure formed by asSL1 could be sufficient to inhibit the association of U2AF1. Alternatively, Ku binding to asSL1 could physically obstruct the association of this critical splicing factor. Our DMS-experiments demonstrate that asSL1 unfolds more frequently in the absence of Ku, which is consistent with Ku directly binding to this RNA structure in cells to stabilize it. Future experiments will be necessary to determine whether the stabilization of asSL1 or direct competition between the Ku complex and the splicing machinery are responsible for the repression of cryptic splice acceptors within antisense Alu RNAs. Importantly, the vast majority of intronic antisense Alu elements are not exonized after Ku degradation, suggesting that they are repressed by a distinct mechanism. For example, Ku insensitive antisense Alu element could be efficiently inhibited by hnRNP C or contain particularly stable asSL1 stem loops that do not require Ku binding to strengthen their folding. The evolution of multiple alternative mechanisms to suppress Alu elements from dysregulating RNA splicing highlights the danger they pose to the cells of higher primates. On the other hand, it is likely that it increased transcriptome complexity enabled by Alu element insertions contributed to the evolution of higher primates into such complex organisms.

Importantly, we documented dysregulation of splicing after Ku depletion in four independent cell systems using distinct methodological approaches targeting both Ku subunits. The genes impacted by alternative splicing include factors that participate in essential processes such as translation, ribosome biogenesis, and splicing itself, which likely explains the rapid cell death observed after Ku depletion. It is possible that some of the observed changes in RNA splicing are indirect effects, resulting from expression changes in splicing factors. However, it is important to note that all data included in our analysis stemmed from the earliest available time points, before substantial cell death was observed in U2OS and HCT116 cells, which should minimize secondary effects. The core set of genes aberrantly spliced after Ku depletion in all three cell lines due to antisense Alu elements includes a number of essential genes (e.g. *EIF3B*, *EXOSC9*, *RPL27a*, *PDCD11*). The loss of any of these proteins could in principle result in cell death and due to the broad effect of Ku depletion on splicing it is likely not a single factor that is solely responsible for the observed phenotype. It is possible that innate immune signaling triggered by Alu elements contributes to cell death in HCT116 cells. However, we did not observe activation of this pathway in U2OS or HEK293T cells and in HCT116 cells disrupting innate immune signaling only partially rescues cell viability ^15^. The preference of Ku for antisense Alu elements is consistent with its role in splicing ^15^, considering that only these sequences contain the poly-pyrimidine tracts that are key elements of splice acceptors. In contrast, it is not immediately clear how preferentially binding to antisense Alu elements would prevent dsRNA mediated innate immune signaling triggered by sense Alu sequences.

In total, the results presented here strongly support a model in which DNA-PK (Ku70, Ku80, DNA-PKcs) expression increased in unison with the expansion of Alu elements in the genomes of higher primates to inhibit splice acceptor sites within antisense Alu sequences. We propose that Ku acts as a critical regulator of splicing homeostasis in primate genomes, balancing the pervasive influence of Alu elements on transcript diversity while safeguarding the fidelity of RNA maturation.

## METHODS

### EXPERIMENTAL MODEL AND STUDY PARTICIPANT DETAILS

In this work, we used these human cancer cell lines: U2OS (RRID: CVCL_0042) and HEK293T ((RRID: CRL-3216). In addition, we carried out experiments with Chinese hamster ovary (CHO) cells (RRID: CVCL_4340). All experiments were performed using low-passage cell lines. U2OS cells were obtained from ATCC, and cell line authenticity was routinely verified by monitoring cell morphology via microscopy. All lines were regularly monitored for mycoplasma contamination. Although all cell lines used in this study are of female origin, the gender, sex, and ethnicity were not considered variables in the experimental design or results, as this study investigates fundamental molecular mechanism relevant to all cell types.

## METHOD DETAILS

### Cell Culture

U2OS cells were grown in RPMI 1640 media (Gibco) supplemented with 10% FBS (Gibco), 100 U ml^−1^ penicillin and 100 µg ml^−1^ streptomycin (Gibco) at 37°C and 5% CO_2_. HEK293T and CHO cells were grown in DMEM media (Gibco) supplemented with 10% FBS, 100 U ml^−1^ penicillin and 100 µg ml^−1^ streptomycin at 37°C and 5% CO_2_.

### Construction of HaloTagged cell lines

Halo-Ku70, Halo-XLF, and Halo-DNA-PKcs U2OS cell lines have been described previously^21^. Generation of Halo-Ku80 U2OS and Halo-Ku70 HEK293T cells was accomplished via the same methods. Cells with HaloTags incorporated at the N-terminus were screened first via genomic DNA-PCR and then confirmed by western blot analyses as described previously ^21^.

### Cell viability assay

MTT staining was performed to assess cell viability for 4-day survival assay as well as 96 hours survival time-course of halo-tagged cell lines. Duplicate wells of each cell line were plated into 24-well plates, containing medium with DMSO or 1 µM HaloPROTAC-E (AOBIOUS, CAS: 2365478-58-4). After 4 days of Halo-Protac treatment (or at 24 h, 48 h, 72 h, 96 h time points), cells were treated with 1 mg/ml MTT (GoldBio) solution for 1 h. Medium containing MTT was then removed and 200 µL DMSO was added per well to lyse cells and solubilize the formazan crystals formed. Absorbance was read at 570 nm to determine relative survival. Background reading was collected from blank wells without cells following the same treatment as experiment and control wells.

### Colony forming assay

Clonogenic survival assays were performed in U2OS cells. Briefly, two hundred cells were plated for each transfectant into complete medium containing the indicated dose of calicheamycin in 60 mm diameter tissue culture dishes. After 8 days, cell colonies were stained with 1% (w/v) crystal violet in ethanol to measure relative survival. Four independent experiments were performed, and results were averaged.

### RNA isolation and RT-PCR

Total cellular RNA was isolated using Trizol (Invitrogen) according to the manufacture’s recommendations. cDNA was synthesized using Superscript-RT (ThermoFischer) according to the manufacture’s recommendations. Resultant cDNA was used in RT-PCR experiments using Go-Taq (ThermoFischer). RT-PCR primers are listed in key resource table.

### Western blotting

SDS-PAGE and western transfer were carried out using standard protocols and probed using primary antibodies targeting Ku70 (1:1000, Neomarkers, Ab4), Ku80 (1:1000, XYC), DNA-PKcs (1:1000, 42-27, a generous gift from Tim Carter), phospho-T446 PKR (1:1000, Invitrogen, 111), HRP-FLAG-M2 (1:4000, Sigma Aldrich A8592), and HRP conjugated anti-mouse and anti-rabbit secondary antibodies. For fluorescent HaloTag staining, cells were incubated for 30 minutes with 100 nM of JF549-HaloTag ligand prior to generating cell lysates.

### Gene editing for AluSz deletion

Cas9-targeted gene disruption was performed by cloning duplex oligonucleotides (Integrated DNA Technologies) into pCas2Apuro, generating two gRNAs specific for PAM sites that target the Alu element in intron 9 of EIF3B. HF-Ku70 U2OS cells were transfected with 2 μg each plasmid; 48 hours after transfection, cells were replated at cloning densities in media containing puromycin (1μg/ml). Puromycin was removed after 72 hours. Isolated clones were selected, and screened by PCRs to assess deletion of the Alu element.

### RNA-seq

The Illumina Stranded mRNA Library Prep kit (Illumina, Cat no. 20040534) with IDT for Illumina RNA Unique Dual Index adapters was used for library preparation following the manufacturer’s recommendations but using half-volume reactions. Qubit™ dsDNA HS (ThermoFischer Scientific, Cat no. Q32851) and Agilent 4200 TapeStation HS DNA1000 assays (Agilent, Cat no. 5067-5584) were used to measure quality and quantity of the generated libraries. The libraries were pooled in equimolar amounts, and the Invitrogen Collibri Quantification qPCR kit (Invitrogen, Cat no. A38524100) was used to quantify the pooled library. The pool was loaded onto 2 lanes of a NovaSeq S4 flow cell at Genewiz Sequencing, and sequencing was performed in a 2×150 bp paired end format using a NovaSeq 6000 v1.5 100-cycle reagent kit (Illumina, Cat no. 20028316). Base calling was performed with Illumina Real Time Analysis (RTA; v3.4.4), and the output of RTA was demultiplexed and converted to the FastQ format with Illumina Bcl2fastq ( v2.20.0).

### RNA-seq analysis

RNA sequencing reads were analyzed using the Michigan State University High Performance Computing Center using a custom pipeline built in Snakemake (Version 7.32.4) (https://github.com/kaylaconner/olivelab-rnaseq/tree/main) ^49^. Read quality was assessed using FastQC (Version 0.12.1) ^50^. Read mapping was performed with Bowtie2 (Version 2.5.1) using default settings and the Hg19 genome. Aligned reads were counted using the featureCounts program in Subread (Version 2.0.6) ^51^. Normalized count tables and differential gene expression analysis was done using the DESeq2 package (Version 1.40.2) in R (Version 4.3.0) ^52^. Counts were pre-filtered to only keep genes that had >10 counts in at least three samples. Publicly available RNA-seq data from HCT116 and HEK293 cells were obtained from the Gene Expression Omnibus (GEO) accession GSE294709.

### Alignment and annotation of differentially spliced introns

For all datasets, Fastq files were trimmed with trimgalore (https://github.com/FelixKrueger/TrimGalore) and aligned on GRCh38 with STAR (v2.7.9a), using parameters: --twopassMode Basic --outSAMstrandField intronMotif --outSAMtype BAM SortedByCoordinate --outSAMunmapped Within --outSAMattributes Standard --sjdbGTFfile/home/giovanni/UCSC_TABLES/hg38/gencode.v48.annotation.gtf --sjdbOverhang 100. Differential splicing was quantified using LeafCutter (v0.2.9) with default parameters, which clusters spliced reads based on shared intron–exon boundaries and tests for differential intron excision ratios between experimental and control samples. Aligned RNA-seq reads in BAM format were first processed to extract splice junctions using the leafcutter_cluster_regtools.py utility. Only junctions supported by at least 30 split reads across all samples and detected in ≥3 biological replicates were retained for analysis. Each intron cluster was tested for differential splicing using the Dirichlet-multinomial model implemented in LeafCutter’s leafcutter_ds.R script. Log-likelihood ratio tests (LRT) were used to identify clusters with significant changes in intron usage between knockout and control conditions. P-values were adjusted for multiple testing using the Benjamini– Hochberg false discovery rate (FDR) correction, and clusters with FDR ≤ 0.05 were considered significantly differentially spliced. For each significant cluster, effect sizes were computed as the difference in intron inclusion levels (ΔPSI) between conditions. Clusters were further classified as Up or Down based on the sign of ΔPSI, indicating increased or decreased intron usage in the knockout condition, respectively. Gene-level associations were determined using GENCODE v48 annotations, assigning each cluster to the host gene(s) overlapping its intronic coordinates. To integrate splicing data with repeat element annotations, introns were intersected with RepeatMasker annotations (GRCh38, UCSC RepeatMasker track) using BEDTools (v2.27.0). This allowed annotation of introns for the presence and orientation of repeat elements relative to the transcriptional strand.

### Gene Ontology analysis

Genes carrying at least one significantly spliced intron cluster (p.adj<0.05 and |logef|>1) were extracted for each dataset, together with the set of genes spliced in all three. Cluster-to-gene assignments were taken from the cluster event classification table and resolved against GENCODE v48, preferring Ensembl gene identifiers and falling back to gene symbols where no identifier was present. Overrepresentation of GO Biological Process terms was tested for each gene set with enrichGO (clusterProfiler) against org.Hs.eg.db, using Ensembl identifiers and the default annotated-genome background. Terms were retained at p<0.05 and q<0.2 after Benjamini–Hochberg correction, with gene set sizes limited to between 10 and 500 genes. Redundant terms were collapsed with simplify at a semantic similarity cutoff of 0.7, and the top 15 remaining terms per set were visualised as dot plots.

### Classification of splicing event types

Significant intron clusters (p.adj<0.05 and |logef|>1) were classified by the configuration of their splice sites. A cluster of three introns sharing one common 5′ donor and one common 3′ acceptor, with the two short introns separated by a gap, was called a single cassette exon (SE-single). Clusters in which all introns shared a common acceptor with multiple donors were called A5SS, and the converse configuration A3SS. Clusters of four or five introns resolvable into two non-overlapping, mutually exclusive cassettes were called MXE-like. Remaining clusters containing at least one cassette triple were called SE-multi, and all others Other_Complex. For downstream summaries, SE-single and SE-multi were merged into SE, and MXE-like and Other_Complex into Other. Each cluster’s splice sites were compared against introns reconstructed from consecutive exons of protein-coding transcripts in GENCODE v48, and clusters were labelled annotated, partial or cryptic according to the fraction of their splice sites present in that reference. Alu elements were taken from the RepeatMasker annotation for hg38, restricted to canonical chromosomes and to the Alu family. Overlap was scored at the junction level, treating each intron as two splice-site positions, and an overlap was called when the position fell within an Alu. Overlap rates were compared per event type against the set of tested but non-significant introns by Fisher’s exact test, and across event types within the significant set by a chi-square test. Alu orientation was assigned relative to the strand of the parental gene, with Alus on the opposite strand called antisense, and the antisense fraction was tested against 0.5 by a binomial test.

For cassette events, all triples of a long skipping intron and two flanking short introns were enumerated within each significant cluster, pairing each upstream short intron with the nearest compatible downstream partner. Triples whose long skipping intron was not itself significant were discarded. Direction was assigned from the sign of the long intron’s ΔPSI: cassettes lost upon depletion were called Ku-activated, and cassettes gained upon depletion Ku-repressed.

### Exon-spanning irAlu enrichment analysis

For each Ku-sensitive cassette exon, the two flanking introns and the cassette body were reconstructed from the four splice-site coordinates of the enumerated triple, following the LeafCutter convention in which an intron start is the last position of the upstream exon and an intron end the first position of the downstream exon. Upstream and downstream were assigned in the direction of transcription according to the strand of the host cluster. Alu elements were taken from the RepeatMasker hg38 annotation, restricted to canonical chromosomes and to the Alu family, and counted per strand within each flanking intron and within the cassette body. A cassette was classified as carrying an exon-spanning irAlu when at least one Alu in the upstream flanking intron and at least one Alu in the downstream flanking intron lay on opposite DNA strands, in either arrangement; all other cassettes were classified as “Other”. Within each dataset, the proportion of Ku-activated to Ku-repressed cassettes was tested against a 50:50 null by a two-sided binomial test for each architectural category, and the two categories were compared by a two-sided Fisher’s exact test on the 2×2 table of direction against architecture, with odds ratios and 95% confidence intervals reported for enrichment of Ku-activated cassettes among exon-spanning irAlu cassettes.

### Correlation between antisense Alu density and splicing magnitude

For each splice junction of Ku-repressed cassette exons, Alu elements from the RepeatMasker hg38 annotation were counted within a symmetric window centered on the junction, separately for each strand, and assigned as sense or antisense relative to the host gene. Windows of 1, 2, 5 and 10 kb were evaluated, with 2 kb taken as the primary window. Counts from the donor and acceptor of an intron were pooled, and local Alu architecture was expressed as log2((antisense+0.5)/(sense+0.5)). The 0.5 pseudocount was used to keep junctions with Alus on only one strand in the analysis with finite values instead of being dropped, since those are the architecturally most extreme cases. Introns with no Alu on either strand within the window were excluded. Introns within a LeafCutter cluster share splice sites and therefore share most of their Alu neighbourhood, so all inference was evaluated against a cluster-level null. Introns were divided into quartiles of the Alu ratio, and the effect was summarized as the ratio of median |ΔPSI| between the antisense-dominated quartile (Q4) and the sense-dominated quartile (Q1). Three estimators were used. First, a cluster-block permutation in which whole per-cluster blocks of Alu ratio were exchanged between clusters of equal size while |ΔPSI| was held fixed, with n=20,000 permutations. Since this null is itself biased above unity, observed folds were assessed against the permuted null rather than against 1. Second, repeated subsampling of one intron per cluster, (n=2,000 draws), with the median fold across draws reported together with its 2.5th and 97.5th percentiles. Third, a rank-based linear mixed model of rank(|ΔPSI|) on the Alu ratio with a random intercept per cluster, fitted with lme4 and lmerTest in R. Spearman correlation between the Alu ratio and |ΔPSI| was assessed under the same permutation scheme. P values were Benjamini– Hochberg adjusted across datasets within each window.

### Fraction of expressed multi-exon genes with Ku-dependent splicing changes

For each dataset, the denominator was defined as the set of GENCODE v48 gene_ids that were both expressed (per-replicate-averaged TPM≥5) and multi-exonic (at least one annotated transcript with n≥2 exons); a protein-coding subset (gene_type == “protein_coding”) was reported in parallel. The numerator was the set of denominator gene_ids harboring at least one significant LeafCutter intron cluster, defined as status == “Success”, cluster p.adj<0.05, and at least one intron with |logef|>1. Each significant cluster was classified into one of four topology categories (SE, A5SS, A3SS, or Other) by a cluster-level classifier operating on the intron coordinate pattern within the cluster (cassette-exon arrangements identified by start/end coincidence with a third intron whose start exceeds the end of the first; A5SS / A3SS by donor-end / acceptor-end coincidence across all cluster introns). A gene was counted in a given event-type bin if at least one of its significant clusters fell into that category.

### Prediction of NMD sensitivity for Ku-REPRESSED cassette exons

Ku-repressed cassettes were tested for predicted introduction of a premature termination codon (PTC) and NMD targeting by the classical 50-nt rule. For each cassette, the parent gene was resolved against the GENCODE v48 annotation, and the per-cassette “principal” transcript was selected as the longest-CDS protein-coding isoform of that gene whose transcribed span contains the cassette body and whose annotated exons do not overlap it. Cassettes whose body fell outside any annotated CDS context were excluded from PTC prediction. The reading frame at the end of the upstream flanking exon of the principal transcript was computed by summing CDS lengths from the start codon to that position. The cassette was then evaluated for two independent PTC-introducing mechanisms: (i) frameshift, defined as cassette body length not divisible by 3; and (ii) in-frame stop, defined as the presence of a TAA/TAG/TGA codon when the cassette body sequence (retrieved from the hg38 genome FASTA via Rsamtools::getSeq, strand-corrected for genes on the minus strand) is translated in the frame established at the upstream flanking exon end. A cassette was classified as “PTC-introducing” if either mechanism applied. The classical 50-nt NMD rule was then applied by computing the distance, in mature-mRNA coordinates, from the predicted PTC position (the first in-frame stop codon for in-frame-stop cassettes; the immediate downstream-exon boundary for frameshift cassettes) to the last exon-exon junction of the mature mRNA with the cassette inserted. Cassettes whose PTC fell ≥50nt upstream of this junction were classified as “NMD_strict” (predicted NMD substrates); the broader “NMD_simple” class includes all PTC-introducing cassettes regardless of distance. Per-cassette predictions were aggregated to the host-gene level and cross-referenced with per-dataset DESeq2 results (p.adj<0.05 and |log2FoldChange|>1) to compute the fraction of downregulated DEGs that overlap a Ku-repressed, NMD-predicted cassette, using two complementary denominators: (a) downregulated DEGs among cassette-affected genes only, and (b) all downregulated DEGs in the dataset.

### Dimethyl Sulfate Chemical Probing

The dimethyl sulfate chemical probing protocol was adapted from Mitchell *et al.*^53^. Control and Halo-PROTAC treated cells were grown for 48 hours to 80% confluency in a 10 cm dish. Media was exchanged for 7.4 ml of fresh media supplemented with 2.1 ml of bicine buffer (pH 8.3) for 3 minutes at room temperature. In cell DMS probing was completed by adding 1.05 ml of 1.7 M dimethyl sulfate in ethanol to the media and incubating at 37°C for 6 minutes. To quench the reaction, 10 ml of 20% beta-mercaptaethanol in PBS was added. Cells are collected by scraping and pelleted for 5 minutes at 4°C at 1000 g.

RNA was extracted from cells by resuspending in 1 mL Trizol for 5 minutes at room temperature, followed by 200 ul of chloroform for 10 minutes. Phases were separated at 12,000 g at 4°C for 10 minutes. RNA was precipitated using one volume of isopropanol, vortexing, and incubating for 10 minutes at room temperature, followed by centrifugation at 12,000 g at 4°C for 8 minutes. The RNA was washed with 1 ml of 75% ethanol and spun at 7500 g for 5 minutes. The ethanol was removed, leaving the pelleted RNA behind. RNA was resuspended in 44 ul of water. Genomic DNA was removed by sing Turbo Dnase (Invitrogen). 5 ul of Turbo buffer was added to the resuspended RNA, followed by 1 ul of Turbo DNase. After incubation at 37°C for 30 minutes, another 1 ul of Turbo DNase was added and incubated again for 30 minutes at 37°C. The RNA was purified using a 1.8x volume of SPRI beads, washing 3x in 70% ethanol, and elution in 30 ul of water.

Relaxed fidelity reverse transcription was carried out using UltraMarathon (RNAConnect): 3 ul dNTPs, 1.5 ul 2 uM gene specific reverse primer, and 15 ul of RNA were incubated at 65°C for 10 minutes, followed by 4°C for 2 minutes. The sample was then supplemented with 10.5 ul of MaP buffer with a final concentration of 50 mM pH 8.3 Tris-HCl, 200 mM KCl, 5 mM DTT, 1 mM MnCl_2_, 20% glycerol and 1 ul of UltraMarathon reverse transcriptase. The reverse transcription was incubated at 42°C for 3 hours, then 95°C for 1 minute.

Prior to library preparation, the cDNA was cleaned up using a G50 column (Cytiva). Step 1 PCR for library preparation was carried out in a 25 ul volume, using 0.02 U/ul Q5 Hot Start Polymerase (NEB), 1.25 ul of each gene specific forward and reverse primer, 1x Q5 reaction buffer, 200 uM dNTPs, and 5 ul of cDNA. Samples were placed into the thermocycler with the following cycles: 98°C for 1 minute; 20 cycles of 98°C for 10 seconds, 66°C for 30 seconds, 72°C for 30 seconds; followed by a final extension of 72°C for 2 minutes. The step 1 PCR product was purified with a 1x ratio of SPRI beads, washing twic with 80% ethanol, and eluting in 15 ul of water. Step 2 PCR was carried out in 50 ul PCR reactions using 2 ng of Step 1 PCR product, 5 ul of each 5 uM indexing primer, 0.02 U/ul Q5 Hot Start Polymerase, 1x Q5 reaction buffer, and 200 uM dNTPs. Step 2 libraries were cycled at 98°C for 1 minute; 10 cycles of 98°C for 10 seconds, 67°C for 30 seconds, 72°C for 30 seconds; and finally, 72°C for 2 minutes. Libraries were cleaned up using 0.8x ratio of SPRI beads, washing twice in 80% ethanol, and eluting in 15 ul of water. Library quality was analyzed using Qubit and Tapestation and were sequenced using 150 x 150 paired-end Illumina next-generation sequencing completed by CD Genomics.

### ShapeMapper 2 DMS mutational profiling analyis

DMS-MaP sequencing data was analyzed using ShapeMapper (version 2.2) in order to calculate mutational frequencies ^54^. ShapeMapper was utilized to process paired-end sequencing FASTQs by performing read quality and primer trimming, aligning reads to the EIF3B reference sequence, and calculating per-nucleotide mutational frequencies. The mutational frequencies of untreated controls were subtracted from the reactivity of DMS treated samples to filter out background mutations and sequencing errors. The mutational frequency is calculated by ShapeMapper based on the number of mutations observed among reads for a specific nucleotide and reactivities were normalized by dividing the mutations frequencies by the mean of the top 10% of mutational frequencies observed in each biological replicate. The fold change in DMS reactivity between control and Halo-PROTAC treated samples was calculated by dividing the normalized reactivity of Halo-PROTAC treated samples by the normalized reactivity of untreated control samples.

## QUANTIFICATION AND STATISTICAL ANALYSIS

All statistical analyses were performed using GraphPad Prism software. For all reported p-values, results were considered non-significant (ns) when p > 0.05. Exact p-values for significant results are provided directly in the figures. Each figure legend includes detailed information on sample size, the number of biological or technical replicates, and the specific statistical analysis used.

## Acknowledgements

We thank our colleague Yu Zhang for many helpful suggestions and careful review of the manuscript. This work was supported by NIH grant (DP2 GM142307) to J. C. S. and by the USDA National Institute of Food and Agriculture (1019208), and Public Health Service (AI048758, AI147634 to K.M.) and the Meek/Hasemann Cancer Research Fund.

## Author contributions

Conceptualization: G.P., J.C.S., and K.M.; Experiments: M.T., M.M., J.C.S., Ga.P., J.R.H., T.J., L.H., K.Y., and K.M. Data Analysis: G.P., Ga.P., A.O., J.G., J.C.S., K.M.; Writing: Original Draft: K.M.; Writing: Review and Editing: G.P., J.C.S., and K.M.

## Data Availability

RNA-seq and DMS-MaP data have been submitted to the gene expression omnibus (GEO, PRJNAXXXX, GSEYYYY). All other data is available from J.C.S. and K.M. upon reasonable request.

## Supplemental Data

**Supplementary table 1.** List of the 72 genes with altered splicing in U2OS, HEK293, and HCT116 cells after depletion of Ku70 (U2OS, HEK293) or Ku80 (HCT116).

---

**Figure S1.**
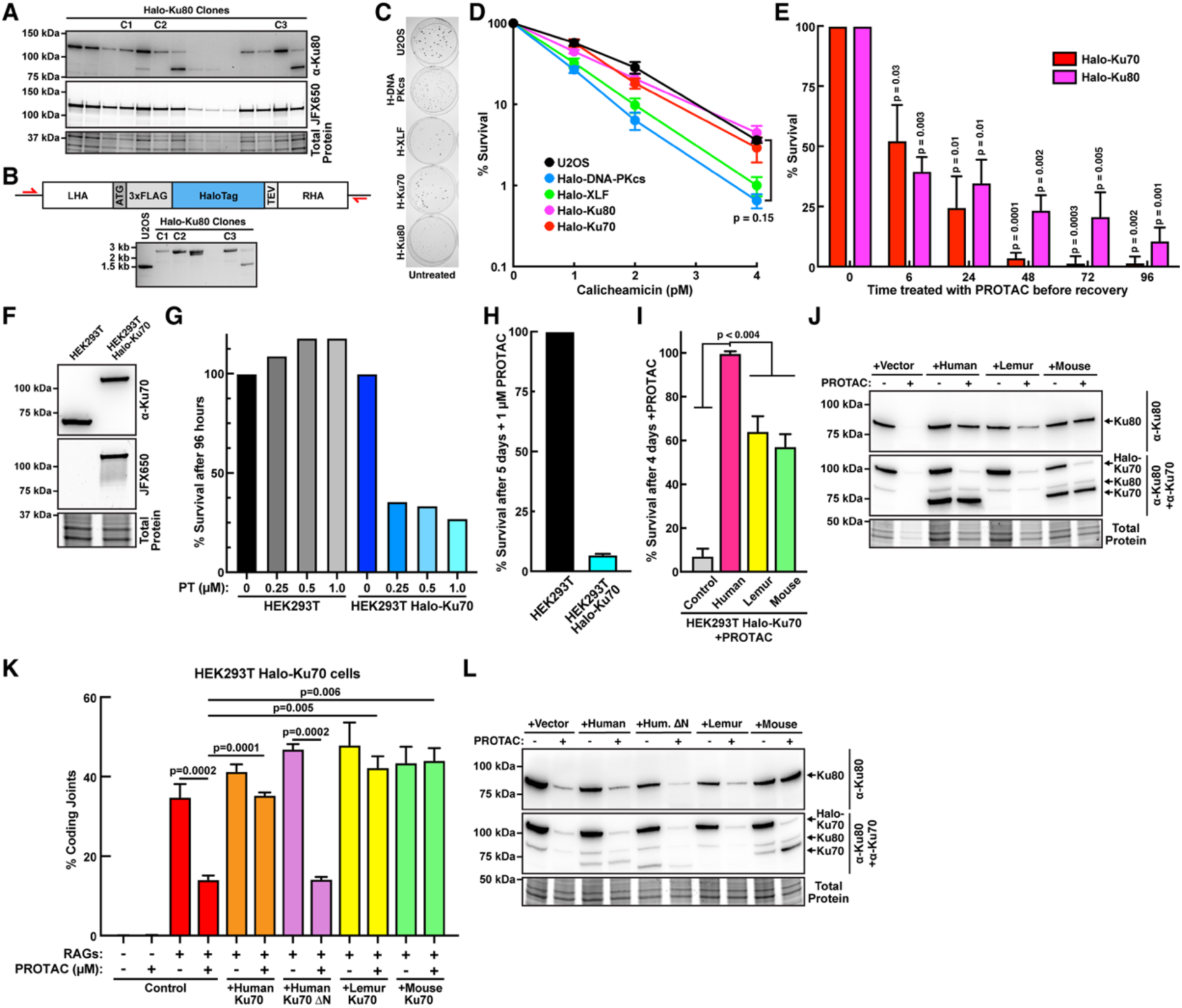
Viability and functional analysis of U2OS cells expressing HaloTagged NHEJ factors and HEK293T cells expressing Halo-Ku70. (**A**) Anti-Ku80 western blot (top), fluorescence gel (middle) and loading control (bottom) of cell lysates generated from U2OS cells expressing Halo-Ku80. (**B**) Cartoon depicting Halo-tag targeting construct and primer positions used to assess targeting as shown in PCR screen (bottom panel). (**C**) U2OS cells were plated at cloning density and stained with crystal violet 8 days later. Colony size is markedly reduced in HF-DNA-PKcs and HF-Ku80 cells. (**D**) U2OS cell strains were plated at cloning densities into complete medium with increasing doses of calicheamicin. Colonies were stained after eight days, and percent survival was calculated. Error bars represent the standard error of the means for six independent experiments (T-test). (**E**) Cell survival of Halo-Ku70 and Halo-Ku80 U2OS cells treated for the indicated time with 1 µM Halo-PROTAC changing to fresh media without the Halo-PROTAC ligand (N=3 biological replicates, Mean±SEM, T-test). (**F**) Western blot for Ku70 (top), fluorescence gel (middle), and total protein staining (bottom) of protein extracts from parental HEK293T and Halo-Ku70 HEK293T cells. (**G**) Cell survival of HEK293T and Halo-Ku70 HEK293T cells in presence of varying concentrations of Halo-PROTAC ligand after 96 hours of exposure. (**H**) Cell survival of HEK293T and Halo-Ku70 HEK293T cells after treatment with 1 µM Halo-PROTAC ligand for 5 days (N=3 biological replicates, Mean±SEM). (**I**) Rescue experiments for Halo-Ku70 expressing HEK293T cells treated with 1 µM Halo-PROTAC for 96 hours, complemented by stable transfection of lemur, mouse, or human Ku70. Cell survival was assessed by MTT staining and compared to blasticidin resistance conferred by the expression plasmids (N=5 biological replicates, Mean±SEM, T-test). (**J**) Western blots probing for Ku70 and Ku80 of Halo-Ku70 HEK293T cells treated with 1 µM Halo-PROTAC, demonstrating that lemur and mouse Ku70 stabilize human Ku80 to confirm expression of Lemur Ku70 and complex formation with human Ku80. (**K**) VDJ recombination assays using transiently transfected reporter and RAG expression plasmids in HEK293T Halo-Ku70 cells stably expressing human wildtype or ΔN Ku70, lemur Ku70, mouse Ku70, or a vector control, in the presence and absence of Halo-PROTAC ligand (N=3 biological replicates, Mean±SEM, T-test). (**L**) Western blots probing for Ku70 and Ku80 of the cells used in the VDJ recombination assay shown in (**K**).

**Figure S2.**
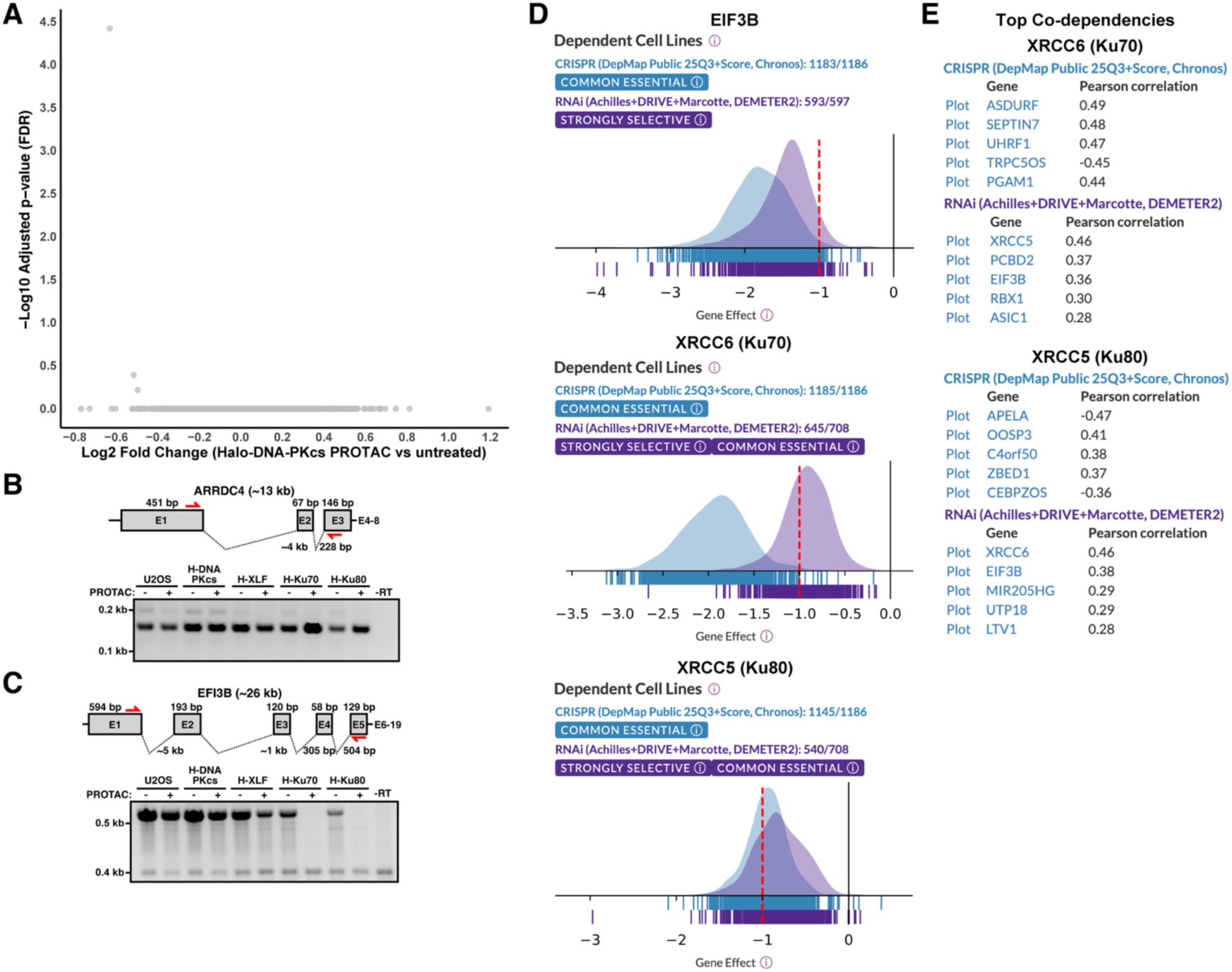
EIF3B is similarly cell essential as Ku70 and Ku80. **(A)** Volcano plot comparing mRNA levels in untreated Halo-DNA-PKcs cells to cells treated with 1 uM Halo-PROTAC for 48 hours. **(B-C)** Validation of upregulation of (B) ARRDC4 or downregulation of (C) EIF3B in Halo-Ku70 cells treated with Halo-PROTAC by RT-PCR. **(D)** The Dependency Map analyses of EIF3B, XRCC6 (Ku70), and XRCC5 (Ku80). All three are similarly essential in over 1000 cell lines as tested by CRISPR knockout experiments or in over 500 cell strains as tested by RNAi experiments. **(E)** Top Dependency Map co-dependencies of Ku70 and Ku80.

**Figure S3.**
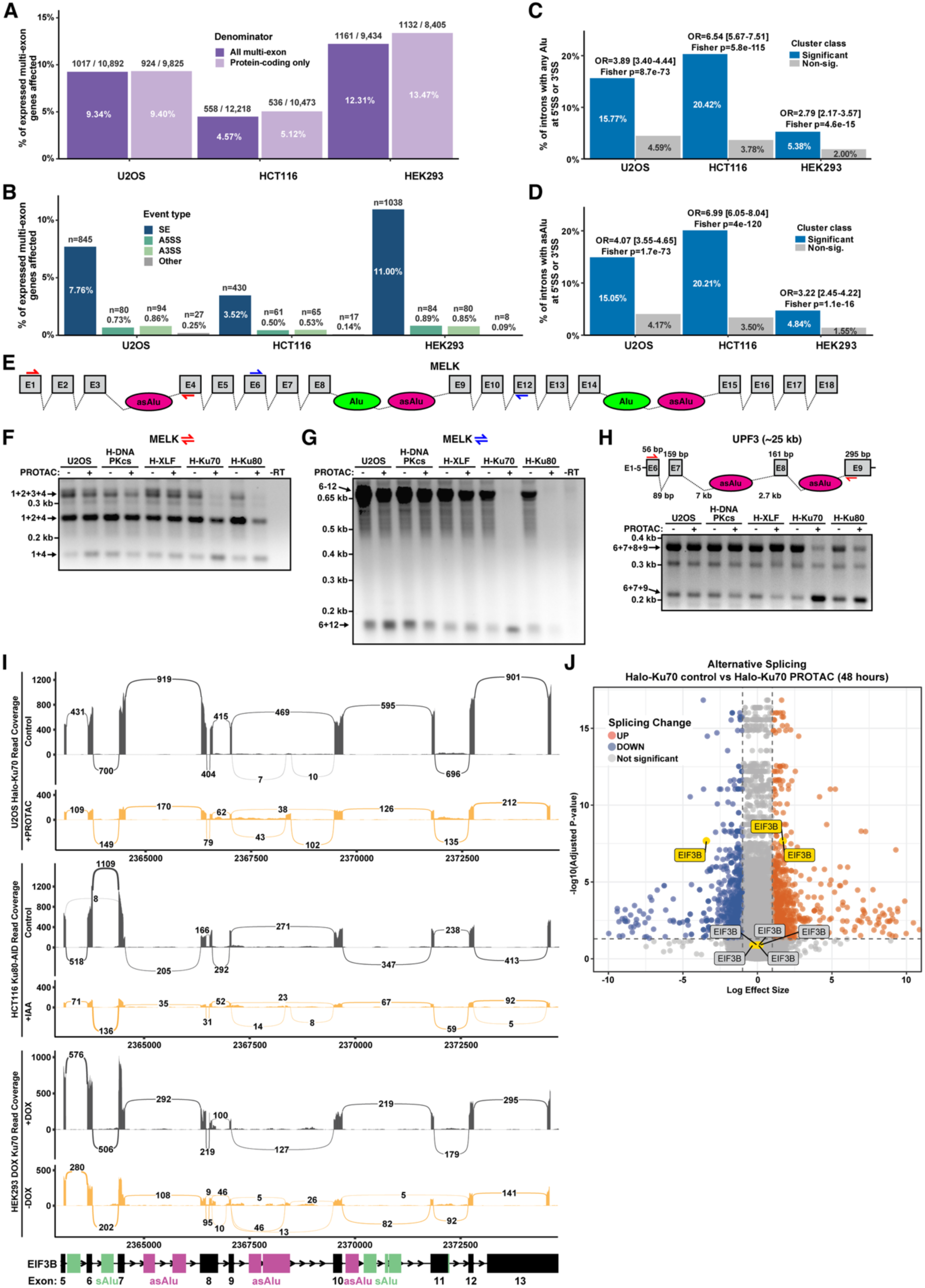
Depletion of Ku causes splicing defects in sequences containing antisense Alu elements. **(A)** Analysis of the fraction of multi-exon or protein coding genes with at least on altered splicing event after Ku70 or Ku80 depletion in U2OS, HCT116, and HEK293 cells. **(B)** Analysis of the fraction of multi-exon genes with specific types of splicing changes after Ku70 or Ku80 depletion in U2OS, HCT116, and HEK293 cells. **(C-D)** Comparison of Alu overlap at splice junctions of significant (p.adj<0.05) and non-significant (p.adj>0.05) alternative splicing events in U2OS, HCT116, and HEK293 cells; **(C)** shows overlap with all Alu elements, **(D)** shows the overlap only with antisense Alu. **(E)** Cartoon depicting the region of MELK transcript that undergoes differential splicing after Ku70 or Ku80 depletion. **(F-G)** RT PCR using the indicated primers to assess splicing of *MELK* between exons 1-4 (**F**) and exons 6-12 (**G**). **(H)** Cartoon and RT-PCR across the first four exons of the UPF3 mRNA in cell lines expressing HaloTagged NHEJ factors treated with Halo-PROTAC ligand. **(I)** Sashimi plots of RNA-seq read coverage and splice junction usage in at the *EIF3B* locus in control (grey) and Halo-PROTAC treated (orange) Halo-Ku70 U2OS cells (top), control (grey) and IAA treated (orange) Ku80-AID cells (middle), and HEK293 DOX Ku70 cells (bottom) in the presence (grey) or absence (orange) of doxycycline. **(J)** Volcano plot showing differential splicing events identified by LeafCutter in U2OS cells after Ku70 depletion, indicating positions of the EIF3B mRNA splice junctions.

**Figure S4.**
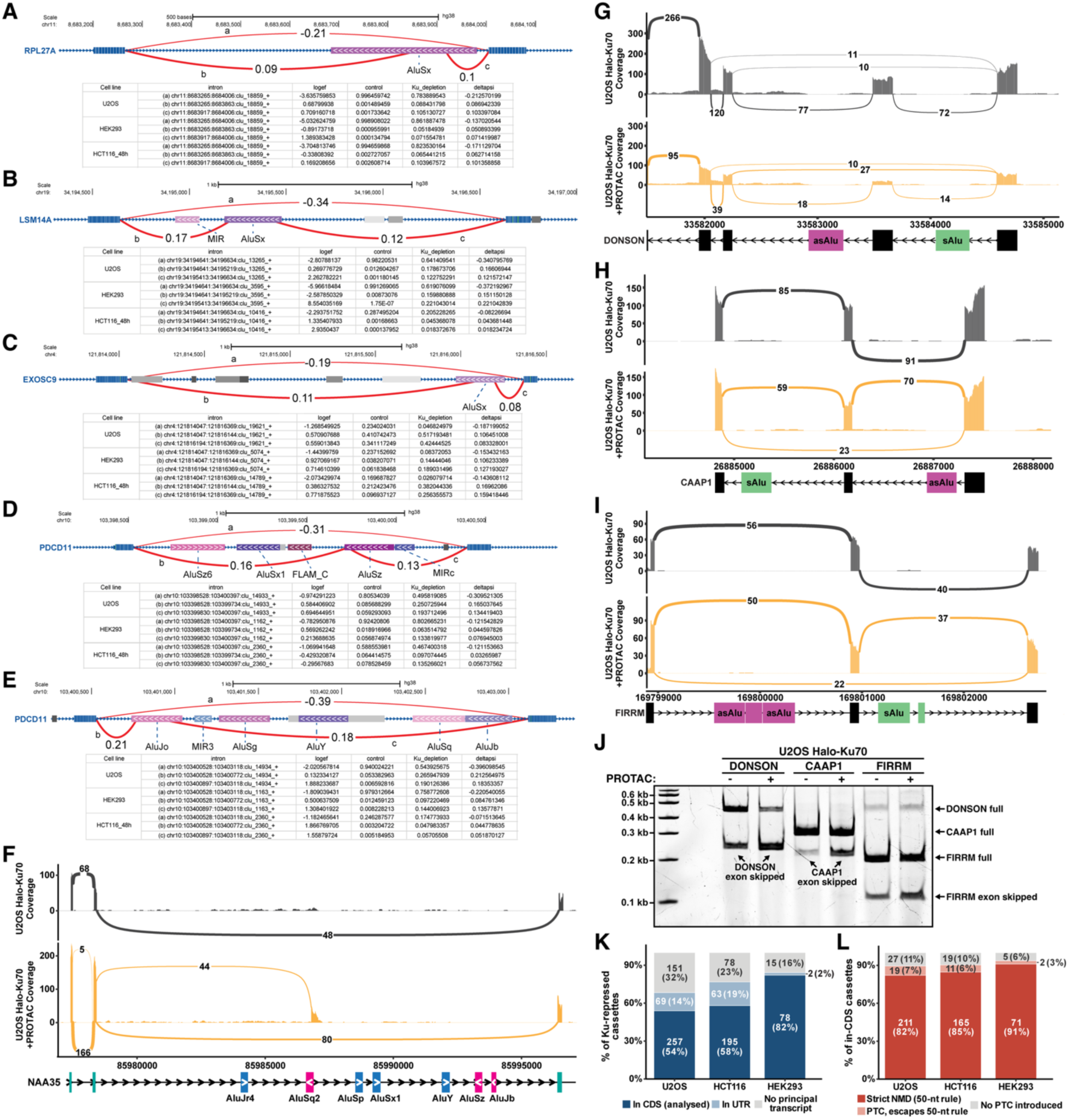
Splicing of antisense Alu element containing sequences is altered after Ku depletion. (A-E), Diagram of splice junctions with start or end points in intronic antisense Alu elements, which are increased after Ku depletion in U2OS, HEK293, and HCT116 cells in the **(A)** *RPL27A,* **(B** *EXOSC9,* **(C)** *LSM14A,* and **(D-E)** *PDCD11* genes. Orientation of the transcripts denoted by blue arrows and orientation of the Alu elements indicted by the white arrows withing the purple shaded boxes. **(F)** Sashimi plot of RNA-seq read coverage and splice junction usage in the *NAA35* gene in control (grey) and Halo-PROTAC treated (orange) Halo-Ku70 U2OS cells. **(G-I)** Sashimi plots of RNA-seq read coverage and splice junction usage in the (G) *CAAP1*, (H) *DONSON*, and (I) *FIRRM* genes in control (grey) and Halo-PROTAC treated (orange) Halo-Ku70 U2OS cells. Orientation of the transcripts denoted by arrows, antisense Alu elements in purple and sense Alu elements in green. **(J)** RT-PCR verification of the exon skipping events in genes containing irAlu pairs shown in (G-I). **(K)** Classification of all Ku-repressed cassette exons per dataset. Each cassette was assigned a principal reference transcript, the longest-CDS protein-coding isoform spanning the cassette in which the cassette is spliced out. Only cassettes within the CDS of that isoform entered PTC prediction; those in a UTR or without an assignable isoform are shown separately. (**L)** Predicted NMD sensitivity of the in-CDS cassettes in (K). Inclusion was scored for a PTC arising from a frameshift (cassette length not a multiple of three) or an in-frame stop in the cassette body, then tested against the 50-nt rule. Counts and percentages are shown within segments.

**Figure S5.**
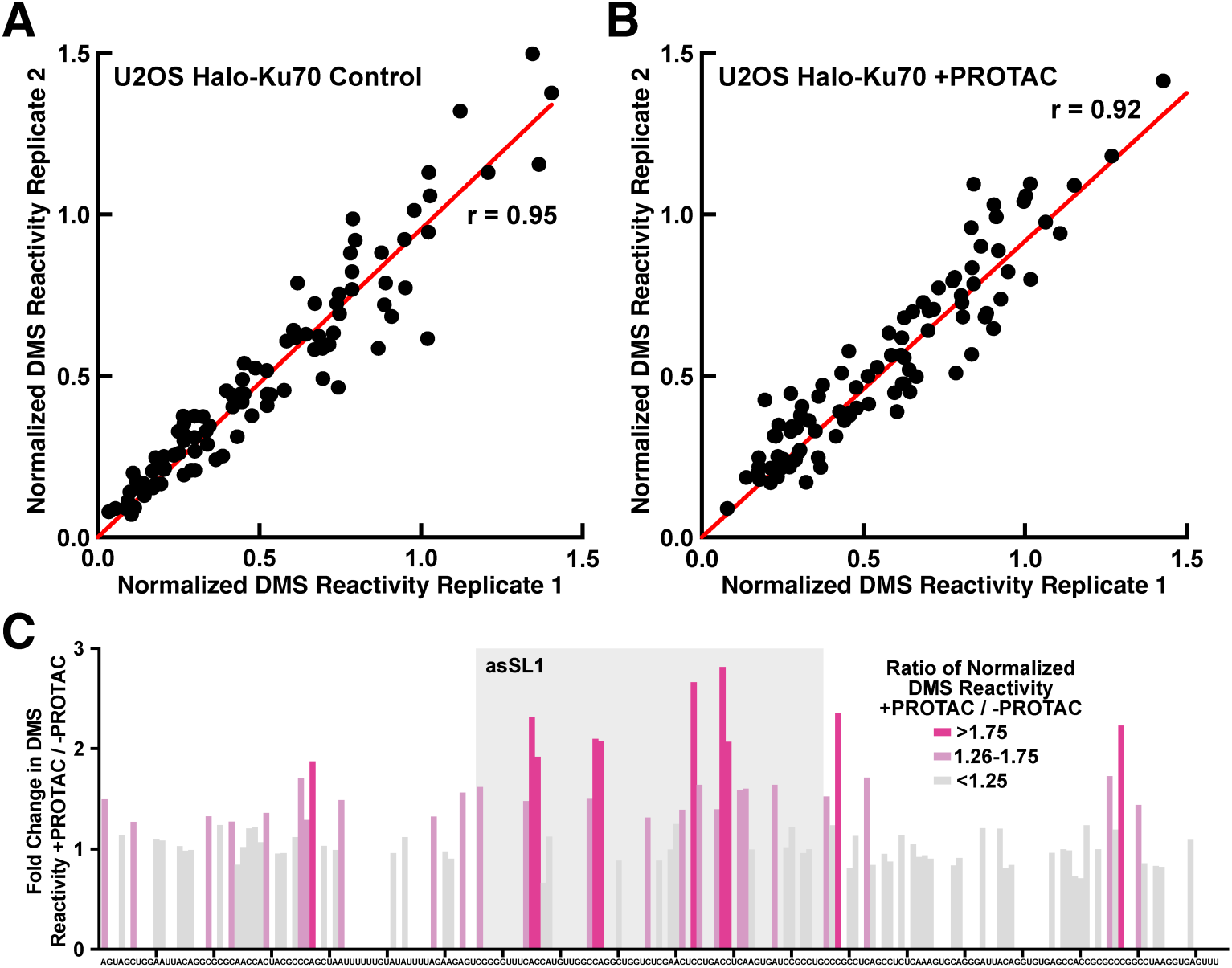
DMS-MaP analysis of asSL1. **(A)** PCR analysis of the deletion of the AluSz element from intron 9 of the EIF3B gene with primers distal to the sgRNA sequences used. **(A-B)** Scatter plots comparing the normalized DMS reactivity of experimental replicates of **(A)** control and **(B)** Halo-PROTAC treated Halo-Ku70 U2OS cells (r = Pearson coefficient). **(C)** Fold change of normalized DMS reactivity for the RNA amplicon from the antisense AluSz element in intron 9 of the *EIF3B* gene spanning the asSL1 stem loop of Halo-PROTAC treated Halo-Ku70 U2OS cells relative to untreated controls. Grey box indicates the asSL1 sequence.

## References

1. Meek, K., Gupta, S., Ramsden, D.A., and Lees-Miller, S.P. (2004). The DNA-dependent protein kinase: the director at the end. Immunological reviews 200, 132–141. 10.1111/j.0105-2896.2004.00162.x.

2. Buehl, C.J., Goff, N.J., Hardwick, S.W., Gellert, M., Blundell, T.L., Yang, W., Chaplin, A.K., and Meek, K. (2023). Two distinct long-range synaptic complexes promote different aspects of end processing prior to repair of DNA breaks by non-homologous end joining. Molecular cell. 10.1016/j.molcel.2023.01.012.

3. Goff, N.J., Mikhova, M., Schmidt, J.C., and Meek, K. (2024). DNA-PK: A synopsis beyond synapsis. DNA repair 141, 103716. 10.1016/j.dnarep.2024.103716.

4. Chaplin, A.K., Hardwick, S.W., Stavridi, A.K., Buehl, C.J., Goff, N.J., Ropars, V., Liang, S., De Oliveira, T.M., Chirgadze, D.Y., Meek, K., et al. (2021). Cryo-EM of NHEJ supercomplexes provides insights into DNA repair. Molecular cell 81, 3400–3409 e3403. 10.1016/j.molcel.2021.07.005.

5. Meek, K., Kienker, L., Dallas, C., Wang, W., Dark, M.J., Venta, P.J., Huie, M.L., Hirschhorn, R., and Bell, T. (2001). SCID in Jack Russell terriers: a new animal model of DNA-PKcs deficiency. Journal of immunology 167, 2142–2150.

6. Finnie, N.J., Gottlieb, T.M., Blunt, T., Jeggo, P.A., and Jackson, S.P. (1995). DNA-dependent protein kinase activity is absent in xrs-6 cells: implications for site-specific recombination and DNA double-strand break repair. Proceedings of the National Academy of Sciences of the United States of America 92, 320–324.

7. Pascarella, G., Conner, K.N., Goff, N.J., Carninci, P., Olive, A.J., and Meek, K. (2024). Compared to other NHEJ factors, DNA-PK protein and RNA levels are markedly increased in all higher primates, but not in prosimians or other mammals. DNA repair 142, 103737. 10.1016/j.dnarep.2024.103737.

8. Pascarella, G., Hon, C.C., Hashimoto, K., Busch, A., Luginbuhl, J., Parr, C., Hin Yip, W., Abe, K., Kratz, A., Bonetti, A., et al. (2022). Recombination of repeat elements generates somatic complexity in human genomes. Cell 185, 3025–3040 e3026. 10.1016/j.cell.2022.06.032.

9. Stankiewicz, P., and Lupski, J.R. (2002). Genome architecture, rearrangements and genomic disorders. Trends in genetics: TIG 18, 74–82. 10.1016/s0168-9525(02)02592-1.

10. Song, X., Beck, C.R., Du, R., Campbell, I.M., Coban-Akdemir, Z., Gu, S., Breman, A.M., Stankiewicz, P., Ira, G., Shaw, C.A., and Lupski, J.R. (2018). Predicting human genes susceptible to genomic instability associated with Alu/Alu-mediated rearrangements. Genome research 28, 1228–1242. 10.1101/gr.229401.117.

11. Li, G., Nelsen, C., and Hendrickson, E.A. (2002). Ku86 is essential in human somatic cells. Proceedings of the National Academy of Sciences of the United States of America 99, 832–837. 10.1073/pnas.022649699.

12. Fattah, F.J., Lichter, N.F., Fattah, K.R., Oh, S., and Hendrickson, E.A. (2008). Ku70, an essential gene, modulates the frequency of rAAV-mediated gene targeting in human somatic cells. Proceedings of the National Academy of Sciences of the United States of America 105, 8703–8708. 10.1073/pnas.0712060105.

13. Woodbine, L., Neal, J.A., Sasi, N.K., Shimada, M., Deem, K., Coleman, H., Dobyns, W.B., Ogi, T., Meek, K., Davies, E.G., and Jeggo, P.A. (2013). PRKDC mutations in a SCID patient with profound neurological abnormalities. The Journal of clinical investigation 123, 2969–2980. 10.1172/JCI67349.

14. Kelly, R.D., Parmar, G., Bayat, L., Maitland, M.E.R., Lajoie, G.A., Edgell, D.R., and Schild-Poulter, C. (2023). Noncanonical functions of Ku may underlie essentiality in human cells. Sci Rep 13, 12162. 10.1038/s41598-023-39166-7.

15. Zhu, Y., Li, A., Maji, S., Lee, B.J., Korn, S.M., Gertie, J.A., Dorrity, T.J., Wang, J., Wang, K.J., Pelletier, A., et al. (2025). Ku limits RNA-induced innate immunity to allow Alu expansion in primates. Nature 643, 562–571. 10.1038/s41586-025-09104-w.

16. Wang, Y., Ghosh, G., and Hendrickson, E.A. (2009). Ku86 represses lethal telomere deletion events in human somatic cells. Proceedings of the National Academy of Sciences of the United States of America 106, 12430–12435. 10.1073/pnas.0903362106.

17. Chen, L.L., and Yang, L. (2017). ALUternative Regulation for Gene Expression. Trends in cell biology 27, 480–490. 10.1016/j.tcb.2017.01.002.

18. Zarnack, K., Konig, J., Tajnik, M., Martincorena, I., Eustermann, S., Stevant, I., Reyes, A., Anders, S., Luscombe, N.M., and Ule, J. (2013). Direct competition between hnRNP C and U2AF65 protects the transcriptome from the exonization of Alu elements. Cell 152, 453–466. 10.1016/j.cell.2012.12.023.

19. Deininger, P. (2011). Alu elements: know the SINEs. Genome Biol 12, 236. 10.1186/gb-2011-12-12-236.

20. Aktas, T., Avsar Ilik, I., Maticzka, D., Bhardwaj, V., Pessoa Rodrigues, C., Mittler, G., Manke, T., Backofen, R., and Akhtar, A. (2017). DHX9 suppresses RNA processing defects originating from the Alu invasion of the human genome. Nature 544, 115–119. 10.1038/nature21715.

21. Mikhova, M., Goff, N.J., Janovic, T., Heyza, J.R., Meek, K., and Schmidt, J.C. (2024). Single-molecule imaging reveals the kinetics of non-homologous end-joining in living cells. Nature communications 15, 10159. 10.1038/s41467-024-54545-y.

22. Caine, E.A., Mahan, S.D., Johnson, R.L., Nieman, A.N., Lam, N., Warren, C.R., Riching, K.M., Urh, M., and Daniels, D.L. (2020). Targeted Protein Degradation Phenotypic Studies Using HaloTag CRISPR/Cas9 Endogenous Tagging Coupled with HaloPROTAC3. Curr Protoc Pharmacol 91, e81. 10.1002/cpph.81.

23. Ponicsan, S.L., Kugel, J.F., and Goodrich, J.A. (2010). Genomic gems: SINE RNAs regulate mRNA production. Current opinion in genetics & development 20, 149–155. 10.1016/j.gde.2010.01.004.

24. Walters, R.D., Kugel, J.F., and Goodrich, J.A. (2009). InvAluable junk: the cellular impact and function of Alu and B2 RNAs. IUBMB Life 61, 831–837. 10.1002/iub.227.

25. Hasler, J., and Strub, K. (2006). Alu elements as regulators of gene expression. Nucleic acids research 34, 5491–5497. 10.1093/nar/gkl706.

26. Nakama, M., Otsuka, H., Ago, Y., Sasai, H., Abdelkreem, E., Aoyama, Y., and Fukao, T. (2018). Intronic antisense Alu elements have a negative splicing effect on the inclusion of adjacent downstream exons. Gene 664, 84–89. 10.1016/j.gene.2018.04.064.

27. Nakama, M., Imanaka, B., and Kimoto, Y. (2025). Intrinsic Alu affects for RNA splicing in a minigene model. Biochem Biophys Rep 42, 102002. 10.1016/j.bbrep.2025.102002.

28. Daniel, C., Behm, M., and Ohman, M. (2015). The role of Alu elements in the cis-regulation of RNA processing. Cellular and molecular life sciences: CMLS 72, 4063–4076. 10.1007/s00018-015-1990-3.

29. Shao, Z., Flynn, R.A., Crowe, J.L., Zhu, Y., Liang, J., Jiang, W., Aryan, F., Aoude, P., Bertozzi, C.R., Estes, V.M., et al. (2020). DNA-PKcs has KU-dependent function in rRNA processing and haematopoiesis. Nature 579, 291–296. 10.1038/s41586-020-2041-2.

30. Li, Y.I., Knowles, D.A., Humphrey, J., Barbeira, A.N., Dickinson, S.P., Im, H.K., and Pritchard, J.K. (2018). Annotation-free quantification of RNA splicing using LeafCutter. Nature genetics 50, 151–158. 10.1038/s41588-017-0004-9.

31. Holmes, M.E., and Hertel, K.J. (2026). Interdependent Regulation of Alternative Splicing by Serine/Arginine-Rich and Heterogeneous Nuclear Ribonucleoprotein Splicing Factors. Genes (Basel) 17. 10.3390/genes17010078.

32. Wickramasinghe, V.O., Gonzalez-Porta, M., Perera, D., Bartolozzi, A.R., Sibley, C.R., Hallegger, M., Ule, J., Marioni, J.C., and Venkitaraman, A.R. (2015). Regulation of constitutive and alternative mRNA splicing across the human transcriptome by PRPF8 is determined by 5’ splice site strength. Genome Biol 16, 201. 10.1186/s13059-015-0749-3.

33. Kreahling, J., and Graveley, B.R. (2004). The origins and implications of Aluternative splicing. Trends in genetics: TIG 20, 1–4. 10.1016/j.tig.2003.11.001.

34. Sela, N., Mersch, B., Gal-Mark, N., Lev-Maor, G., Hotz-Wagenblatt, A., and Ast, G. (2007). Comparative analysis of transposed element insertion within human and mouse genomes reveals Alu’s unique role in shaping the human transcriptome. Genome biology 8, R127. 10.1186/gb-2007-8-6-r127.

35. Lee, H., Cao, X., Parada, G.E., Li, J.D., Iosub, I.A., Braunschweig, U., Rebelo, K., Ule, J., Wan, Y., and Blencowe, B.J. (2026). Alu-mediated RNA duplexes are associated with widespread exon skipping across primate transcriptomes. Genome biology 27. 10.1186/s13059-026-04029-7.

36. Zubradt, M., Gupta, P., Persad, S., Lambowitz, A.M., Weissman, J.S., and Rouskin, S. (2017). DMS-MaPseq for genome-wide or targeted RNA structure probing in vivo. Nature methods 14, 75–82. 10.1038/nmeth.4057.

37. Wang, J., Weatheritt, R., and Voineagu, I. (2022). Alu-minating the Mechanisms Underlying Primate Cortex Evolution. Biol Psychiatry 92, 760–771. 10.1016/j.biopsych.2022.04.021.

38. Xia, B., Zhang, W., Zhao, G., Zhang, X., Bai, J., Brosh, R., Wudzinska, A., Huang, E., Ashe, H., Ellis, G., et al. (2024). On the genetic basis of tail-loss evolution in humans and apes. Nature 626, 1042–1048. 10.1038/s41586-024-07095-8.

39. Sen, S.K., Han, K., Wang, J., Lee, J., Wang, H., Callinan, P.A., Dyer, M., Cordaux, R., Liang, P., and Batzer, M.A. (2006). Human genomic deletions mediated by recombination between Alu elements. American journal of human genetics 79, 41–53. 10.1086/504600.

40. Hedges, D.J., and Deininger, P.L. (2007). Inviting instability: Transposable elements, double-strand breaks, and the maintenance of genome integrity. Mutation research 616, 46–59. 10.1016/j.mrfmmm.2006.11.021.

41. Ade, C., Roy-Engel, A.M., and Deininger, P.L. (2013). Alu elements: an intrinsic source of human genome instability. Curr Opin Virol 3, 639–645. 10.1016/j.coviro.2013.09.002.

42. Adelmant, G., Calkins, A.S., Garg, B.K., Card, J.D., Askenazi, M., Miron, A., Sobhian, B., Zhang, Y., Nakatani, Y., Silver, P.A., et al. (2012). DNA ends alter the molecular composition and localization of Ku multicomponent complexes. Molecular & cellular proteomics: MCP 11, 411–421. 10.1074/mcp.M111.013581.

43. Dvir, A., Peterson, S.R., Knuth, M.W., Lu, H., and Dynan, W.S. (1992). Ku autoantigen is the regulatory component of a template-associated protein kinase that phosphorylates RNA polymerase II. Proceedings of the National Academy of Sciences of the United States of America 89, 11920–11924.

44. Yoo, S., and Dynan, W.S. (1998). Characterization of the RNA binding properties of Ku protein. Biochemistry 37, 1336–1343. 10.1021/bi972100w.

45. Jurica, M.S., and Moore, M.J. (2003). Pre-mRNA splicing: awash in a sea of proteins. Molecular cell 12, 5–14. 10.1016/s1097-2765(03)00270-3.

46. Jurica, M.S., Licklider, L.J., Gygi, S.R., Grigorieff, N., and Moore, M.J. (2002). Purification and characterization of native spliceosomes suitable for three-dimensional structural analysis. RNA 8, 426–439. 10.1017/s1355838202021088.

47. Wilkinson, M.E., Charenton, C., and Nagai, K. (2020). RNA Splicing by the Spliceosome. Annual review of biochemistry 89, 359–388. 10.1146/annurev-biochem-091719-064225.

48. Yoshida, H., Park, S.Y., Sakashita, G., Nariai, Y., Kuwasako, K., Muto, Y., Urano, T., and Obayashi, E. (2020). Elucidation of the aberrant 3’ splice site selection by cancer-associated mutations on the U2AF1. Nature communications 11, 4744. 10.1038/s41467-020-18559-6.

49. Koster, J., and Rahmann, S. (2018). Snakemake-a scalable bioinformatics workflow engine. Bioinformatics 34, 3600. 10.1093/bioinformatics/bty350.

50. andrews, S. (2010). FastQC: A quality control tool for high throughput sequence data. Available from: http://www.bioinformatics.babraham.ac.uk/projects/fastqc/.

51. Liao, Y., Smyth, G.K., and Shi, W. (2014). featureCounts: an efficient general purpose program for assigning sequence reads to genomic features. Bioinformatics 30, 923–930. 10.1093/bioinformatics/btt656.

52. Love, M.I., Huber, W., and Anders, S. (2014). Moderated estimation of fold change and dispersion for RNA-seq data with DESeq2. Genome Biol 15, 550. 10.1186/s13059-014-0550-8.

53. Mitchell, D., Cotter, J., Saleem, I., and Mustoe, A.M. (2023). Mutation signature filtering enables high-fidelity RNA structure probing at all four nucleobases with DMS. Nucleic Acids Res 51, 8744–8757. 10.1093/nar/gkad522.

54. Busan, S., and Weeks, K.M. (2018). Accurate detection of chemical modifications in RNA by mutational profiling (MaP) with ShapeMapper 2. RNA 24, 143–148. 10.1261/rna.061945.117.

